# Protein induced membrane phase transition facilitates leishmania infection

**DOI:** 10.1101/2020.03.08.982454

**Authors:** Achinta Sannigrahi, Junaid Jibran Jawed, Subrata Majumdar, Syamal Roy, Sanat Karmakar, Krishnananda Chattopadhyay

**Author notes:** Structural Biology & Bio-Informatics Division, CSIR-Indian Institute of Chemical Biology, 4, Raja S. C. Mallick Road, Kolkata 700032, India.

## Abstract

Although host membrane is known to play critical roles in the internalization of leishmania parasites inside macrophages (Mϕ), any detailed mechanistic understanding is missing. We show here that KMP-11, a small immunogenic protein of *Leishmania Donovani* (LD) facilitates the infection process by binding to Mϕ membrane through its N-terminal domain (1-19AA). This binding results in a membrane phase transition that occurs at a threshold protein/lipid ratio, which is linked to the change in membrane tension. KMP-11 induced phase transition is also associated with lipid raft disruption and T-cell deactivation. Finally, using a combination of tryptophan-scanning mutagenesis and synthesized peptides, we develop a mathematical exposition, which demonstrates that hydrophobic moment (μ_H_) and the number of residues involved in a mirror sequence (N) at the interacting N-terminal are governing factors for the membrane phase transition, which facilitates infection process.

## Introduction

The term leishmaniasis encompasses multiple clinical syndromes –the cutaneous, mucosal and visceral forms, which are caused by different members of leishmania species. Regardless of diseases manifestation, leishmania parasites are obligatory intracellular pathogens and macrophage (Mϕ) is indispensable for parasite survival, replication, and differentiation. Such obligate intracellular parasite employs a variety of mechanisms to invade host cells and then to evade immune response giving rise to a spectrum of health disorders of the host.^1^ The entry process of the parasites into the Mϕ is complex, which requires careful investigation to understand microbial pathogenesis. One of the major structural components of the surface membrane of leishmania parasites is kinetoplastid membrane protein-11 (KMP-11), which has been implicated in the regulation of the overall lipid bilayer pressure of the parasite membrane.^2–3^ In *Leishmania Donovani* (LD), KMP-11 remains tightly associated with LPG and is present essentially in equimolar amount with LPG.^4^ This KMP-11-LPG complex decreases as a function of parasite subculture coupled with the decrease in parasite virulence.^5^

We found that the following two observations with KMP-11 are important. First, we (and others) have seen significant sequence homology between KMP-11, bovine apolipoprotein A-I and apolipoprotein A-IV.^6^ In addition, a comparison between the thermodynamic properties of KMP-11 and protein members belonging to apolipoprotein family suggest that KMP-11 could be considered as an additional entry in this group of proteins.^7^ Interestingly, a link between the presence of cholesterol in host membrane and the extent of infection has been suggested, which is validated by epidemiological data, which show that the population with less exposure to cholesterol is more likely to be affected by the infection.^8^ Using these, we speculated that the parasite LD uses an apolipoprotein like protein KMP-11 to transport cholesterol from the host membrane to facilitate productive infection. Second, we have recently observed KMP-11 induced pore formation, a property this protein may use to facilitate the entry process of the parasite.^9^ From these two results, we hypothesize that KMP-11 binding to host membrane may have profound implications in the disease mechanism.

To validate this hypothesis, we studied the interactions between KMP-11 and membrane to obtain a mechanistic understanding of how this regulates the disease mechanism. We found that the presence of KMP-11 facilitated and presumably mimicked LD infection, which we established by measuring not only the infection propensity but also associated parameters, like IL2 expression and lipid rafts dislocation. We then showed that KMP-11 modulated LD infection through a novel phase transition mechanism in the lipid membrane, in which the lipid molecules changed their fluidity as monitored by different fluorescence and SAXS assays. In order to understand which region of the protein is responsible for this lipid phase transition, we prepared a number of single tryptophan mutants, in which a tryptophan residue was placed in different parts of the protein. We showed that a sequence stretch at the N-terminal region of KMP-11 (amino acids, 1-19) is responsible for the membrane binding and phase transition. We subsequently prepared different synthetic peptides based on the N-terminal sequence (1-19) to establish that this region generates a Hydrophobic Moment, which plays critical roles in the phase transition. A mathematical model was then developed, in which the Hydrophobic Moment and the number of amino acid residues (N) forming the mirror sequence of the interacting domain at the N terminal, was found to be responsible for all three events, namely membrane binding, the phase transition of the lipid, and the process of infection. The biophysical results using synthetic membranes were validated by measuring directly the infection propensity in Mϕ cell lines.

## Results

### KMP-11 mimics LD infection

We first investigated the effect of LD infection on membrane fluidity (MF). This was done using two popular assays: by measuring a) fluorescence anisotropy (FA) using 1,6-diphenylhexatriene (DPH) and b) generalized polarization (GP) using laurdan. Both DPH and laurdan are commonly used membrane sensitive probes. It has been established that an increase in MF results in the decrease in FA and GP.^10^ Both assays (FA, Figure 1a and GP, Figure 1b) showed significant increase in MF in 12 h infected Mϕ (LD-Mϕ). Both assays also showed that the treatment of cholesterol partially offset the effect of LD infection (Figure 1a, b). Interestingly, we found that KMP-11 mimicked the effect of LD. Both FA (Figure 1a, inset) and GP (Figure 1b, inset) showed a dose dependent decrease (and an increase in MF) when 2h KMP-11 treated normal Mϕ (KMP-Mϕ) was used (Figure S4a,b). In this condition also, the addition of cholesterol partially reversed the change in MF (Figure 1a,b). We noted that the nature of the dose dependence of FA (inset, Figure 1a) was different from that of GP (inset, Figure 1b), the former being hyperbolic while the later sigmoidal. Although the reasons are unknown to us, this difference may happen as a result of the difference in how probe molecules (DPH vs laurdan) would interact with and penetrate the membrane. Subsequently, we compared the T-cell stimulating ability between KMP-Mϕ and normal macrophages (Mϕ) (Figure 2a). Using I-A^d^ restricted anti-LACK T-cells hybridoma and subsequent IL2 production in the presence or absence of LACK antigen, we observed that co-culture of anti-LACK-T-cells with Mϕ without LACK antigen failed to produce IL2 (Figure 2a). In the similar condition, the addition of LACK antigen showed significant IL2 production. In contrast, IL2 production from anti LACK T-cells decreased as the dose of KMP-11 increased for pulsing the Mϕs, which were used as antigen presenting cells.

**Figure 1:**
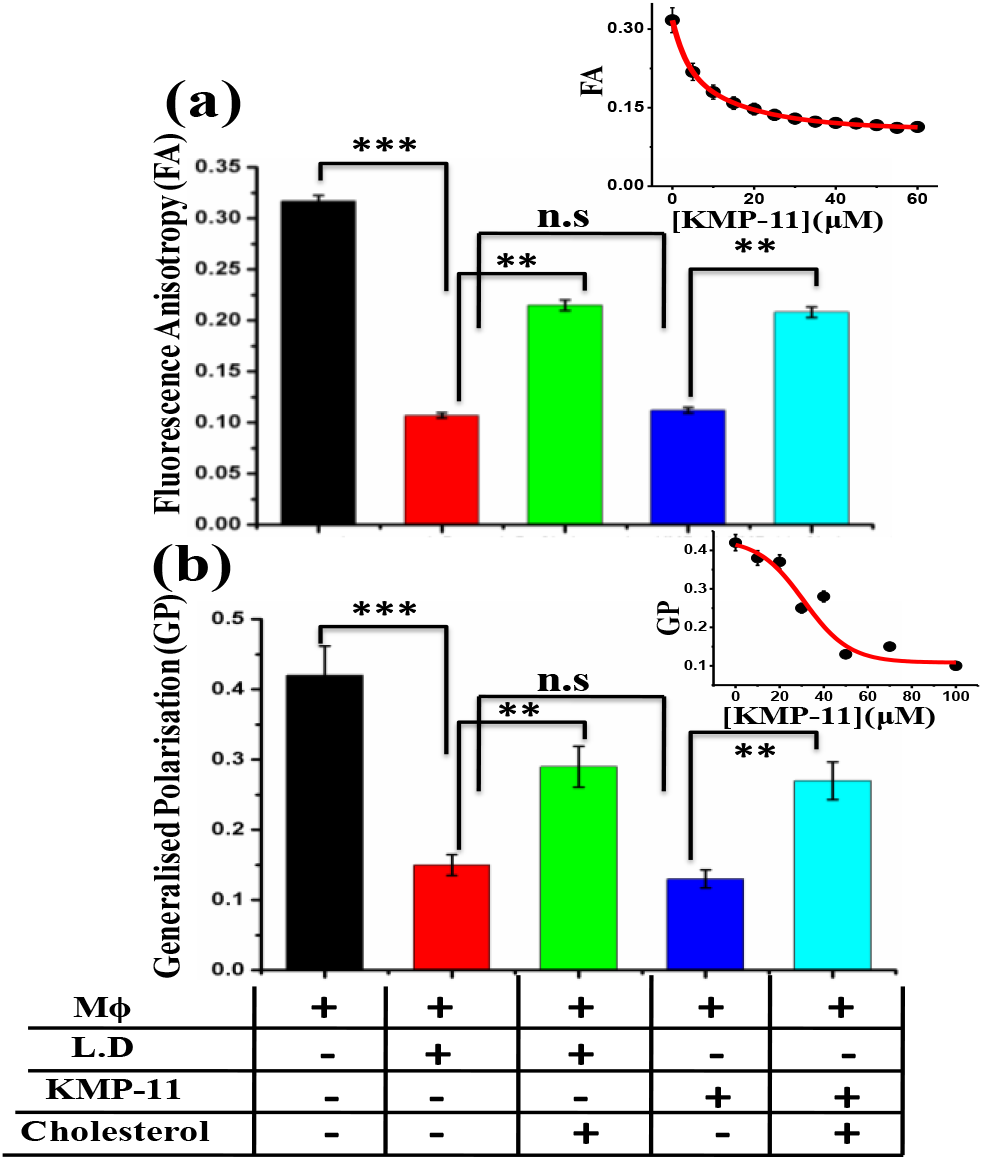
Determination of membrane fluidity of Mϕ membrane under different treatment conditions. (a) FA of Mϕ membrane under the treatment of LD, KMP-11 and Cholesterol. These FA values were determined using DPH. Typical protein concentration was 50 μM and the liposomal cholesterol concentration was taken 100 μM for each experiment (in liposomal cholesterol Lipid/cholesterol ratio was taken 1:2). ** is considered to be significant (p value is <0.01) and *** is considered highly significant (p value is < 0.001) whereas n.s denoted as non-significant. Inset showed the change in FA of Mϕ with increasing concentration of KMP-11.The solid line through the data points was obtained from the hyperbolic fit. (b) GP of laurdan fluorescence of Mϕ membranes under the treatment of LD, KMP-11 and Cholesterol. Typical protein concentration used was 50 μM and the liposomal cholesterol concentration was 100 μM for each experiment (in liposomal cholesterol Lipid/cholesterol ratio was taken 1:2). ** is considered to be significant (p value is <0.01) and *** is considered as highly significant (p value is < 0.001) whereas n.s denoted as nonsignificant. Inset showed the change in GP of Mϕ with increasing concentration of KMP-11.The solid line through the data points was obtained from the sigmoidal fit. + and − signs denoted the presence and absence of different entities.

**Figure 2:**
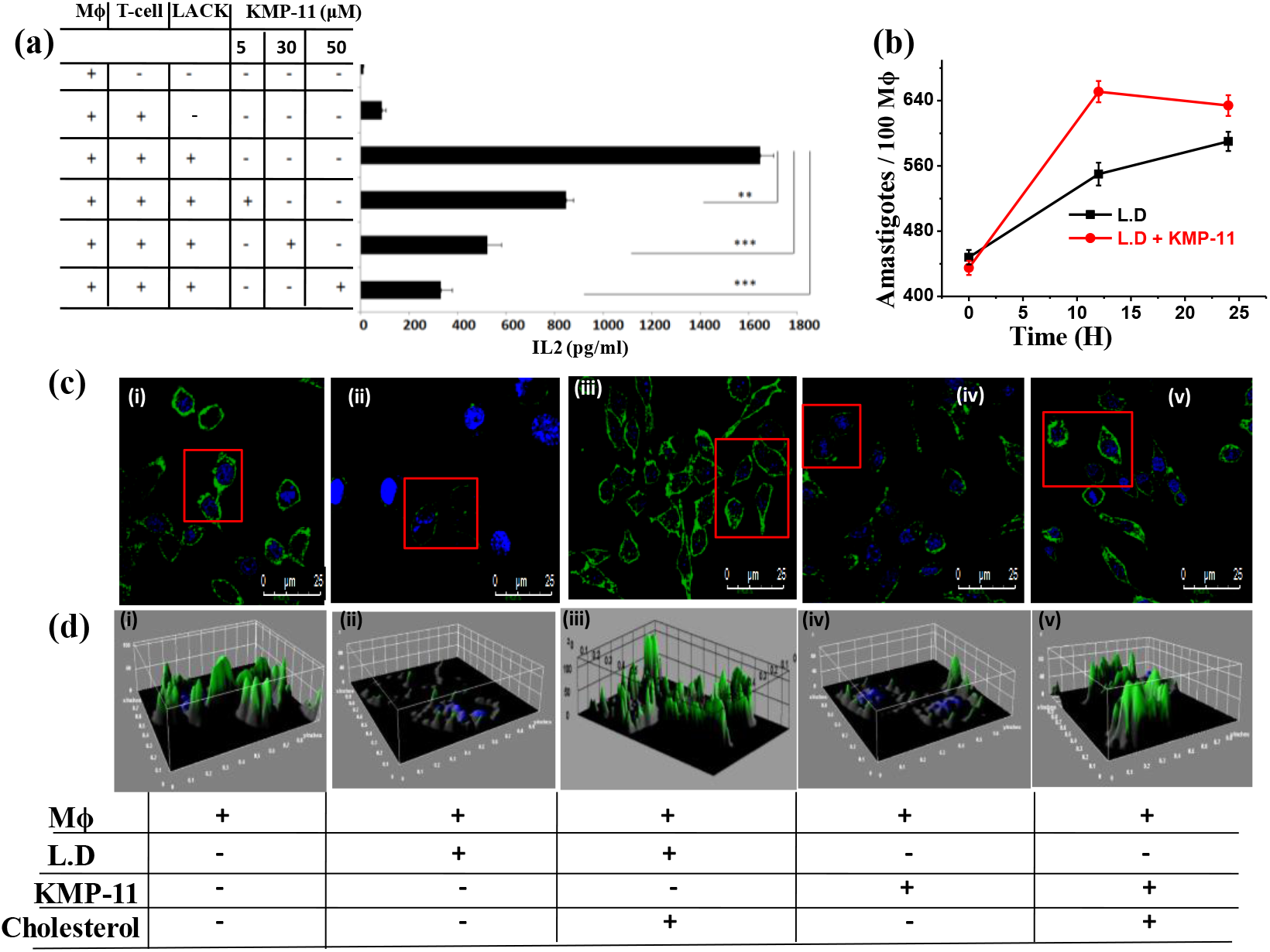
KMP-11 induced phase transition driven subsequent effects in Mϕ cells in terms of IL2 expression, infection propensity and lipid raft dislocation. (a) IL2 expression level under different treatments. Here 5 μM concentration was used as low dose of KMP-11 and 30 μM as medium dose while 50 μM was used as high dose. The Mϕs were cultured with T cell hybridoma – LMR7.5 (T cell) which gets activated by LACK antigen. ** is considered to be significant (p value is <0.01) and *** is considered as highly significant (p value is < 0.001). (b) Parasite load inside Mϕ in a time frame from the treatment of KMP-11. (c) Confocal images of Mϕ rafts and decrease in raft intensity under treatment of KMP-11 and LD and the recovery of the raft after cholesterol treatment. CTX-B-FITC was used as raft marker. Images of (i) control Mϕ, (ii) LD-Mϕ, (iii) LD-Mϕ with cholesterol, (iv) KMP-Mϕ and (v) KMP-Mϕ with cholesterol have been mentioned. (d) 3D interactive plots on the basis of raft’s fluorescence intensities designated the raft morphology of (i) control Mϕ, (ii) LD-Mϕ,(iii) LD-Mϕ with cholesterol, (iv) KMP-Mϕ and (v) KMP-Mϕ with cholesterol. The green color corresponds to the raft marker whereas the blue color denotes the nucleus as stained by DAPI. + and − signs denoted the presence and absence of respective species.

Next, we infected the Mϕs with stationary phase LD parasites for 12 h and then allowed the intracellular parasites to grow. We considered 12 hours LD infection as our time zero (0 hour) of post-infection. The intracellular parasites were counted per100 Mϕs. At 0 hour, the number of intracellular parasites was 448, which increased to 530 at 12 hours (Figure 2b). Interestingly, when we used KMP-11 pulsing of Mϕ and subsequent LD infection, this value was increased to 650 at 12 hours (Figure 2b). These data suggested that the presence of KMP-11 facilitated LD entry rate into Mϕ.

Since it has already been shown that that LD affects raft assembly,^11^ we wanted to find out if KMP-11 causes similar effects. We investigated the binding of Cholera toxin-B (CTX-B)-FITC to cell surface as a marker of raft in control Mϕs (Figure 2c(i)), LD-Mϕs (Figure 2c(ii)), and KMP-Mϕs (Figure 2c(iv)). The binding of CTX-B expressed as 3D interactive plots is shown in Figure 2d (i-v).We observed from the confocal images that CTX-B binds to normal Mϕs (Figure 2c(i) and 2d(i)). However, under LD infection (Figure 2c(ii) and 2d(ii)) and in the presence of KMP-11 (Figure 2c(iv) and 2d(iv)), there was a significant decrease in CTX-B binding to the cell surface which was restored by cholesterol liposome treatment (Figure 2c(iii) and 2d(iii) for LD-Mϕ and Figure 2c(v) and 2d(v) for KMP-11-Mϕ respectively).

### KMP-11 binds to membranes, which is inhibited by cholesterol

In the previous section, we established that KMP-11 mimics LD infection, affecting Mϕ raft assembly and decreasing IL2 production. We also observed that the addition of cholesterol offsets the effect of KMP-11. Subsequently, we wanted to understand the molecular details of how KMP-11 does that. For most of the experiments discussed in this section, we used model membranes made of DPPC (DPPC SUVs). We chose DPPC because it is the major component of mammalian Mϕ membrane.^12^ In addition, membrane fluidity assays using KMP-Mϕ showed similar behaviors of FA and GP values in both Mϕ membrane and DPPC SUVs (Figure 3a).

**Figure 3:**
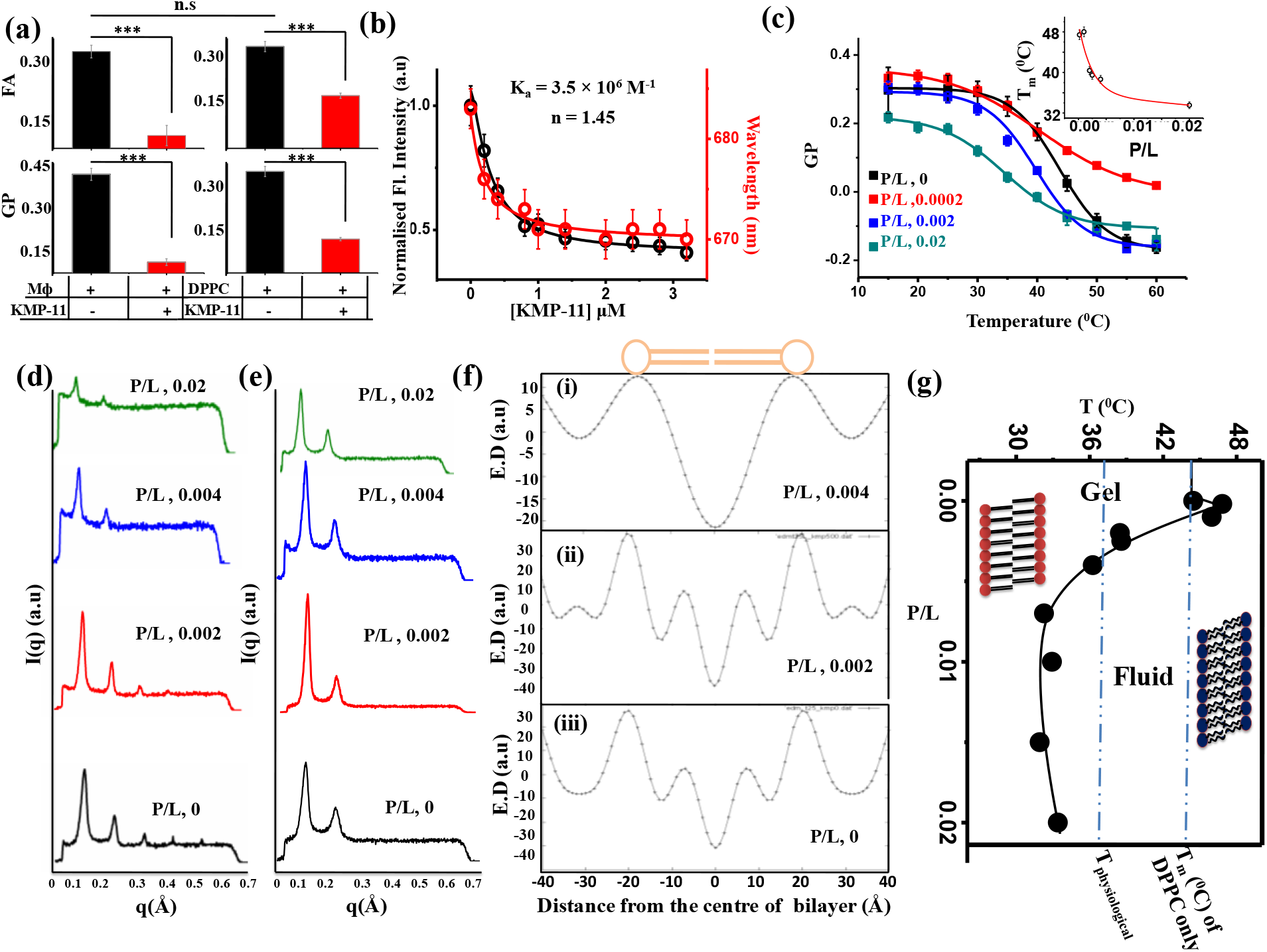
Binding and phase transition in phospholipid model membrane by KMP-11. (a) FA and GP values of DPPC model membrane and Mϕ membrane in absence and presence of KMP-11.*** stands for the change which was highly significant whereas n.s denoted nonsignificant changes.(b) Binding of KMP-11 with DPPC model membranes was monitored by decrease of DiI C-18 fluorescence intensity (left y axis) as well as the blue shifts of the fluorophore (right y axis) with increasing concentration of added proteins. (c) The variation in GP with increase in temperature at different P/L molar ratios. Inset shows the change in T_m_ with P/L.SAXS data of DPPC MLV at different P/L ratios at (d) 25 °C and (e) 50 °C. (f) Electron Density Maps (EDM) of DPPC MLV at three different P/L ratios:(i) 0.004, (ii) 0.002 and (iii) 0.(g) A typical phase diagram of KMP-11 induced DPPC phase transition. T_physiological_ indicates 37°C.

Once we established DPPC as a convenient model for Mϕ, we performed a membrane binding assay of KMP-11 using a membrane specific fluorophore DiI C-18 (Figure 3b, Figure S5a, Table S2). KMP-11 showed one order of magnitude stronger binding towards saturated phospholipid DPPC (*K*_*a*_ = 3.5×10^6^ M^−1^) when compared with unsaturated phospholipid DOPC binding (*K*_*a*_ = 2.7 × 10^5^ M^−1^).^13^ We also found out that the addition of cholesterol resulted in significant decrease in binding affinity (Figure S5b).

### KMP-11 binding changes the properties of lipid molecules and induces a phase transition in membrane

We then carried out a series of experiments to determine if the binding between KMP-11 and membrane results in any change in the properties of the protein and/or lipids. Using far UV CD (which monitors protein secondary structure), we determined that the binding of DPPC did not result in any significant change in the secondary structure (FigureS5c) of KMP-11. In contrast, we monitored the properties of the lipids using FT-IR. The change at the choline moiety (at the head groups) was monitored by –C-N-C-vibrations (Figure S6a), while PO_2_ vibrations were used for the phosphotidyl group (Figure S6b). In addition, the change in the tail region of the lipid was monitored by symmetric stretching (Figure S6c) and symmetric deformation of CH_2_ (Figure S6d) bond vibrations. A schematic diagram of the lipid indicating different regions is shown in Figure S6. We observed significant change both at the head and tail regions of the lipid, which were induced by protein binding (Figure S6a-d, black vs. red curves).

Detailed analyses of FT-IR data provided additional insights into the protein binding induced change in properties of the lipid. For example, the bands at 1728 cm^−1^ and 1742cm^−1^ represent the hydrogen bonded and non-hydrogen bonded stretching frequencies respectively. A comparison of these bands in the absence and presence of KMP-11 clearly suggested an increase in the extent of non-hydrogen bonded ester carbonyl vibration (Figure S6e and S6f). It has been reported earlier that the increase in non-hydrogen bonded carbonyl frequency is responsible for the flexibility as well as mobility and this effect can cause bilayer thinning.^14^ FTIR data thus suggested that the binding of KMP-11created thinner DPPC bilayer with increased flexibility. Further, we used the CH-wagging band wavenumber (1280-1440 cm^−1^) to investigate the conformational changes of the DPPC bilayer due to KMP-11 binding. Vibrational band at 1367 cm^−1^ denotes the nonplanar kink+gtg^/^ conformers (also known as rotamers) of DPPC.^15^ We found an increase in the extent of nonplanar kink+gtg^/^ conformers of DPPC (Figure 4d,e,f), which occurred as a result of KMP-11 binding.

**Figure 4:**
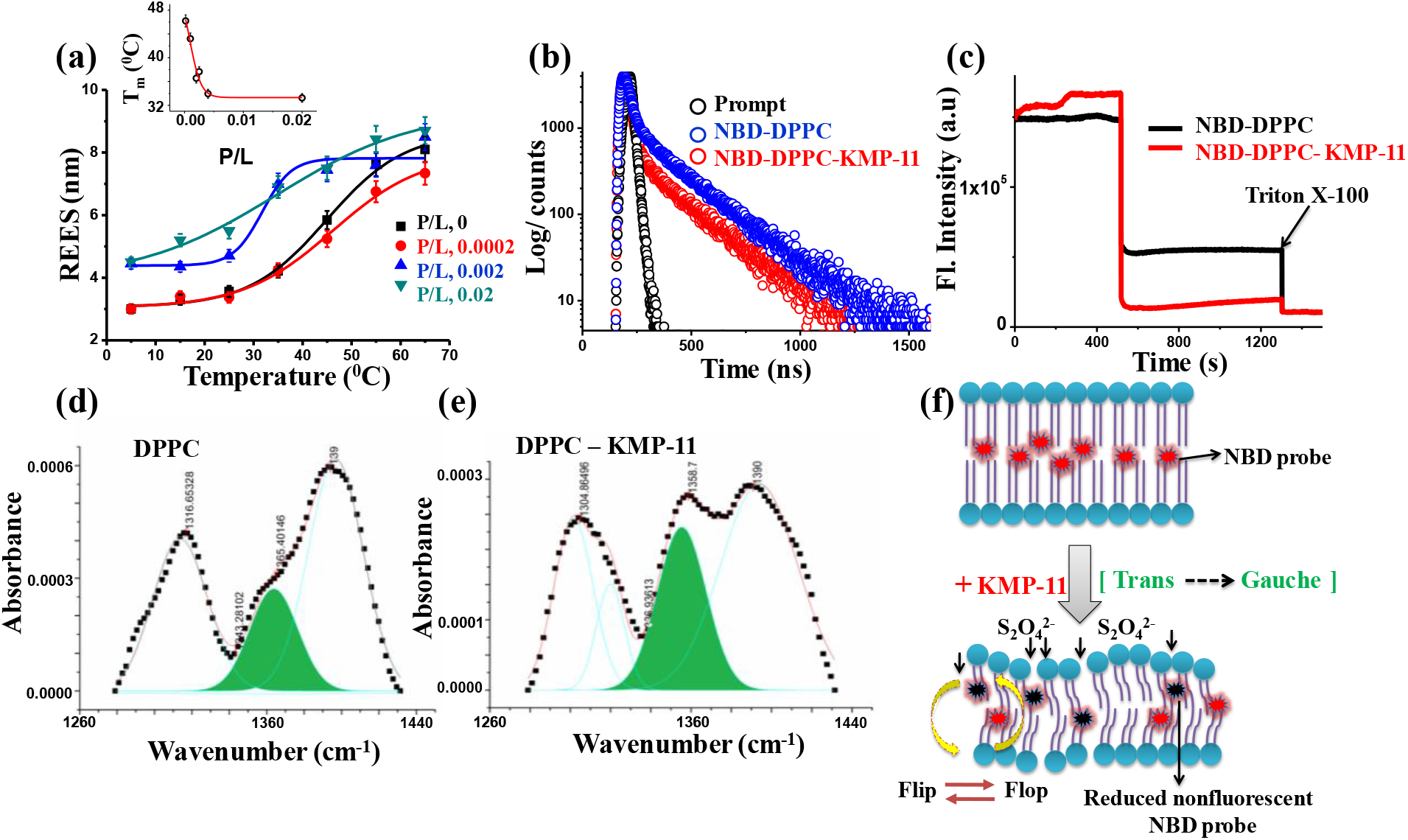
Measurements of the governing factors behind KMP-11 induced phase transition. (a) REES of DPPC-NBD PC at different P/L ratio against temperature. Inset shows the plot of chain melting temperature against P/L. (b) Fluorescence lifetime of DPPC-NBD-PC in absence and presence of KMP-11. (c) Flip-flop dynamics of NBD-PC in presence and absence of KMP-11.FTIR spectral signatures of the CH_2_ wagging band frequency of DPPC in absence (d) and presence (e) of KMP-11. This FTIR data suggested the increase of gauche rotamers (indicated by the green regions) of DPPC due to protein binding.

Since the binding of protein changes the properties of the lipid, it was important to determine if this change at the lipid molecule translates into a change at the membrane level. To understand this, we first determined gel (relatively rigid lipid) to fluid (more flexible) phase transition temperature of DPPC membrane (T_m_) using the temperature dependent red shift of laurdan fluorescence emission (Figure 3c, Figure S7, Table S3). Inset of Figure 3c showed the variation of T_m_ with KMP-11/DPPC (P/L) ratios. We observed that the increase in the relative concentration of the protein in P/L reduced T_m_ (Figure 3c, inset, Table S3). At P/L of 0.004 and above, DPPC remained in its fluid phase even at the physiological temperature (which is much lower than the chain melting temperature of DPPC in the absence of protein, 44°C). We used different mole percentage of cholesterol to trace its effect on KMP-11 induced phase transition in DPPC membrane. Addition of cholesterol significantly increased the GP values (Figure S8a) which indicated the restoration of membrane rigidity as well as phase state (gel) by cholesterol. The critical cholesterol/DPPC (C/L) ratio was found to be 0.7, which is sufficient for the membrane gel state recovery (Figure S8b).

We confirmed protein induced phase transition of DPPC using SAXS experiments. Figures 3d and 3e show the intensity profiles obtained from SAXS at different P/L. Measurements were performed at two different temperatures of 25°C and 50°C respectively. A large number of lamellar reflections at 25°C (Figure 3d, bottom panel where P/L=0) and relatively less number of lamellar reflections at 50° C for only DPPC (Figure 3e, bottom panel where P/L=0) characterizes the gel (at 25°C) and fluid (at 50°C) phases respectively. In the presence of KMP-11, the number of lamellar reflections reduced even at 25°C, suggesting the presence of more flexible bilayers with respect to the gel phase (Figure 3d, P/L 0 to 0.02). The SAXS profile at 25 °C for P/L of 0.004 showed the presence of only two distinct reflections, which was identical to the SAXS profile of DPPC fluid phase at 50°C respectively (Figure 3d and Figure 3e, the second panel from the top for both figures). From the SAXS experiments, we inferred that the bilayer undergoes gel to fluid transition at 25°C for P/L of ~ 0.004 and above. We constructed a typical phase diagram of KMP-11 induced phase transition in DPPC membrane suggesting that approximately P/L ~ 0.004 is critical for the successful gel to fluid phase conversion of model membrane at 25°C (Figure 3g).

We complemented above measurements using a number of fluorescence-based assays. The first one was REES (Red Edge Excitation Shift) measurements using NBD PC. REES assay has been used popularly to study membrane phase transition, and its principle is well established.^16^ We found that the value of REES increased with increasing temperature (Figure 4a), from which we calculated T_m_ at varying P/L (the inset, Figure 4a, Figure S10, Table S3). We observed a decrease in T_m_ as we increased the concentration of KMP-11(Figure 4a), indicating slow rate of solvent relaxation relative to the NBD lifetime due to the reorientation of solvent dipoles around the excited state of fluorophore. Such solvent relaxation in the immediate vicinity of fluorophore depends on motional restriction, i.e, the change in the rotational diffusion of lipid molecules induced by protein in the lipid bilayer. We found a decrease in the value of average lifetime (T_avg_) of NBD-PC from 6 ns to 5 ns (Figure 4b) due to membrane-protein interaction, which complemented nicely with the REES data. Next, we carried out dithionite-induced fluorescence quenching experiments using NBD-PC in DPPC membrane, which showed an increase in the relative fraction of NBD analog on the outer leaflet (P_0_) in the presence of KMP-11 (P_0_ for NBD-DPPC~ 64.25% whereas for NBD-DPPC-KMP-11~ 91.89%) (Figure 4c). This result indicated that the protein binding significantly enhanced the flip-flop rate of lipid molecules. This data also implied a reduction of membrane rigidity with concomitant increase in the flexibility.

### Hydrophobic Moment of KMP-11 regulates the phase transition in lipid membranes

Since the interaction between the protein and membrane is expected to be a contributing factor for the membrane phase transition, we determined the region of KMP-11 responsible towards the binding. For this, we first used a computational tool using OPM (Orientation of Proteins in Membrane environment, http://opm.phar.umich.edu/server.php) server. This calculation suggested that the interaction occurs through the N-terminal (residues 1-19) of the protein (Figure S12a). To validate this prediction experimentally, we used tryptophan scanning mutagenesis in which we prepared four single tryptophan mutants (Y5W, Y48W, F62W & Y89W). WT-KMP-11 does not have any tryptophan residue and in these mutations, and hence these mutations (Figure 5a) would enable a small change in the hydrophobicity at this particular stretch where the tryptophan residue was inserted. The difference in binding constant (Δ*K*_*a*_ = *K*_*a,WT*_ − *K*_*a,mutant*_) would provide an estimate of the effect of mutation site (and the small increase in hydrophobicity) on the binding constant. In addition, we calculated the difference in *K*_*SV*_ for each mutants (Δ*K*_*SV*_ = *K*_*SV*_(free)−*K*_*SV*_(membrane bound)) using the fluorescence quenching measurements. We found out that the mutant Y5W (tryptophan inserted at the N-terminal) showed the maximum effect on *ΔK*_*a*_ and *ΔK*_*SV*_ values (Figure 5b inset, Table S2), which clearly validated OPM prediction.

**Figure 5:**
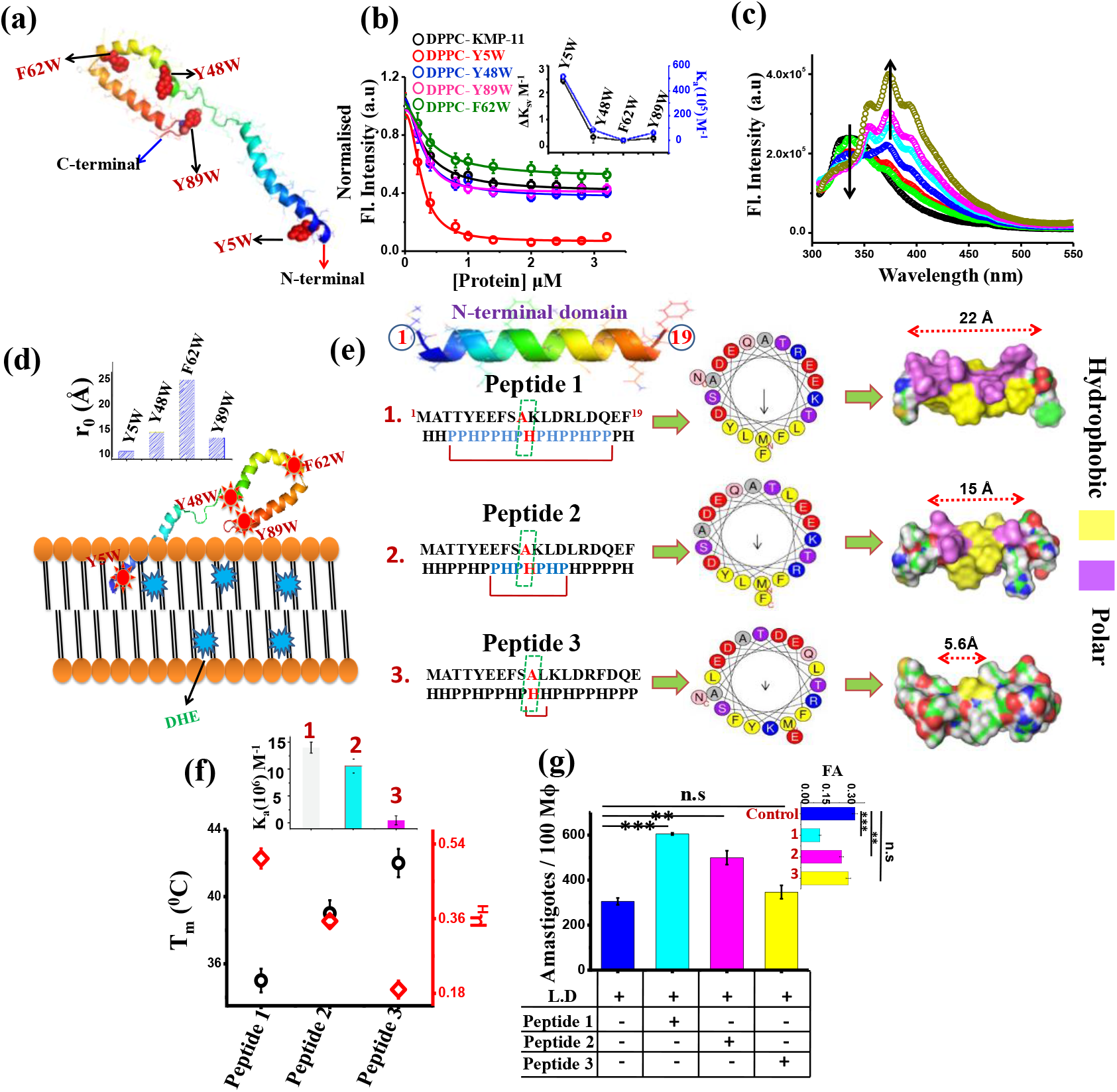
Detection and analysis of the membrane interacting domain of KMP-11. (a) Model structure of KMP-11. The inserted tryptophan residues were marked red. (b) Binding of four single tryptophan mutants with DPPC model membranes as evident from the quenching of DiI fluorescence. Inset showed the change in tryptophan exposure (in terms of *ΔK*_*SV*_) for four single tryptophan mutants as well as their binding affinities (*K*_*a*_, M^−1^) towards model membrane. (c) FRET study of Y5W mutant using the tryptophan residue as donor and membrane bound DHE as acceptor molecule.(d) Positioning of the protein on the membrane as confirmed by the experimental findings. Inset showed the donor acceptor distances as calculated using tryptophan as donor and membrane DHE as acceptor.(e) Sequences of three synthesized peptides possessing different hydrophobic moments as revealed by helical wheel modeling.(f) Hydrophobic moments and phase transition temperatures of DPPC SUV induced by three peptides plotted altogether. Inset showed the plot of binding affinity of three peptides towards model membrane. (g)Parasite load inside macrophages in 12 hrs of incubation with three different peptides.** is considered to be significant (p value is <0.01) and *** is considered as highly significant (p value is < 0.001) whereas n.s denoted as nonsignificant. Inset showed the DPH anisotropy change of macrophage membranes by three peptides.

Penetration of sequence stretch of KMP-11 inside membrane was further determined by calculating energy transfer efficiency (E) between the inserted tryptophan (donor) and membrane bound Dehydroergosterol (DHE, acceptor) using them as FRET pairs (Figure S12a). We observed that Y5W mutant showed highest energy transfer efficiency (E~20%) (Figure5c). In contrast, Y48W and Y89W exhibited intermediate values (E_Y48W_~6.4%; E_Y89W_~8.5%). F62W mutant showed the lowest (E_F62W_~ 1.72%) (Figure S12b-e,Table S4). From the energy transfer results, we found that apparent donor-acceptor distances followed the trend Y5W (11.39Å) < Y89W (13.64Å) < Y48W (14.13Å) < F62W (24.67 Å) (Table S4, Figure 5c-d, Figure S12b-e). The tryptophan scanning strategy hence suggested that hydrophobic interaction plays an important role in the binding, which presumably occurred through the N-terminal (sequence 1-19).

We found that the N-terminal sequence (1-19) (**^1^MATTYEEFSAKLDRLDQEF^19^)** contains a mirror stretch when the sequence was arranged on the basis of residue specific hydrophobicity (H) and polarity (P) (Figure 5e). Since an ideal α–helix makes exactly five turns, we used a helical wheel model to investigate the hydrophobic face and to calculate the hydrophobic moment (μ_H_) of the specific interacting domain. Heliquest compuparam (http://heliquest.ipmc.cnrs.fr/cgi-bin/ComputParamsV2.py) analysis showed that helical wheel model of KMP-11-N-terminal domain (1-19AA) contains a hydrophobic face consisting of **LFMFLY** (Figure 5e, Table S5) with a net hydrophobic moment of 0.505. Subsequently, we designed a few new stretches theoretically using this helical wheel model and altering the amino acids positions so that the hydrophobic moments were gradually altered (Figure 5e, Table S5) without changing the net hydrophobicity. We computed the binding energies between the designed peptides and membranes (ΔG_t_, Kcal/mole), which showed that ΔG_t_ increased with the increase in the μ_H_ of the sequence (Table S5). To investigate in details why sequence symmetry (which makes the difference in hydrophobic moment and membrane affinity) is a critical determinant for the binding phenomenon, we determined the number of residues, which constituted the mirror stretch (N). Table S5 also showed that there was a direct relation between N and ΔG_t_. To validate this relation experimentally, we synthesized three different peptides of varying hydrophobic moments with similar hydrophobicity (namely peptide1, peptide 2 and peptide 3 in Figure 5d). Binding studies of these peptides with DPPC model membrane suggested that binding affinity followed the trend peptide 1> peptide 2> peptide 3 (Figure 5f, inset, Figure S13a). Then, laurdan GP experiments of DPPC SUVs were executed in presence of the peptides. By measuring laurdan fluorescence, we found that of gel-fluid transition temperature (T_m_) was directly proportional to μ_H_ of the peptides (Figure 5e, Figure S13b). Next, we wanted to understand if the change in μ_H_ had the same effect on Mϕ cell lines during LD infection. For this, we measured DPH FA of Mϕ membrane while treated with these peptides. We found that Mϕ membrane fluidity was significantly higher for peptide 1 in comparison to peptide 2. Peptide 3 was found to be inert towards the modulation Mϕ membrane fluidity as evident from the inappreciable change in FA values (Figure 5g inset). Next we studied LD infection propensity in Mϕ cell line under the treatment of three peptides. Interestingly, when we used peptide pulsed Mϕ and subsequent LD infection, intracellular parasite count was found to be noticeably greater for peptide 1 while compared to peptide 2 after 12 hr of incubation (Figure 5g). No considerable increase in parasite load was observed when Mϕ cells were pulsed with peptide 3 (Figure 5g).

## Discussion

We have summarized the presented data using Scheme 1, which aims to explain the roles of KMP-11 in the disease manifestation. In this study, we established that KMP-11 treatment mimics LD infection. The protein itself induces the change in the fluidity of Mϕ membrane, which follows all other subsequent effects (lowering of IL2 expression, increase in parasite load inside Mϕ etc). We also found that, like what happens in LD infection, these KMP-11 induced effects could be reversed by cholesterol treatment. The effect of KMP-11 was triggered by its ability to bind to host membrane inducing a membrane phase transition. Protein-membrane interaction is known to modulate the infection propensity in different disease models but no clear evidence on the protein induced membrane phase transition during pathogenic infection is available in literatures till date. Here we show that KMP-11 binding perturbs the membrane morphology, although the conformation of the protein remains more or less unaltered. Membrane phase transition is associated with a number of changes at the molecular level, which include the increase in flip flop dynamics of the lipid molecules, change in the conformations of the lipid from trans to gauche orientations and bilayer thinning. These morphological alterations are expected to play crucial roles in the lipid raft disruption and T-cell deactivation.

**Scheme 1:**
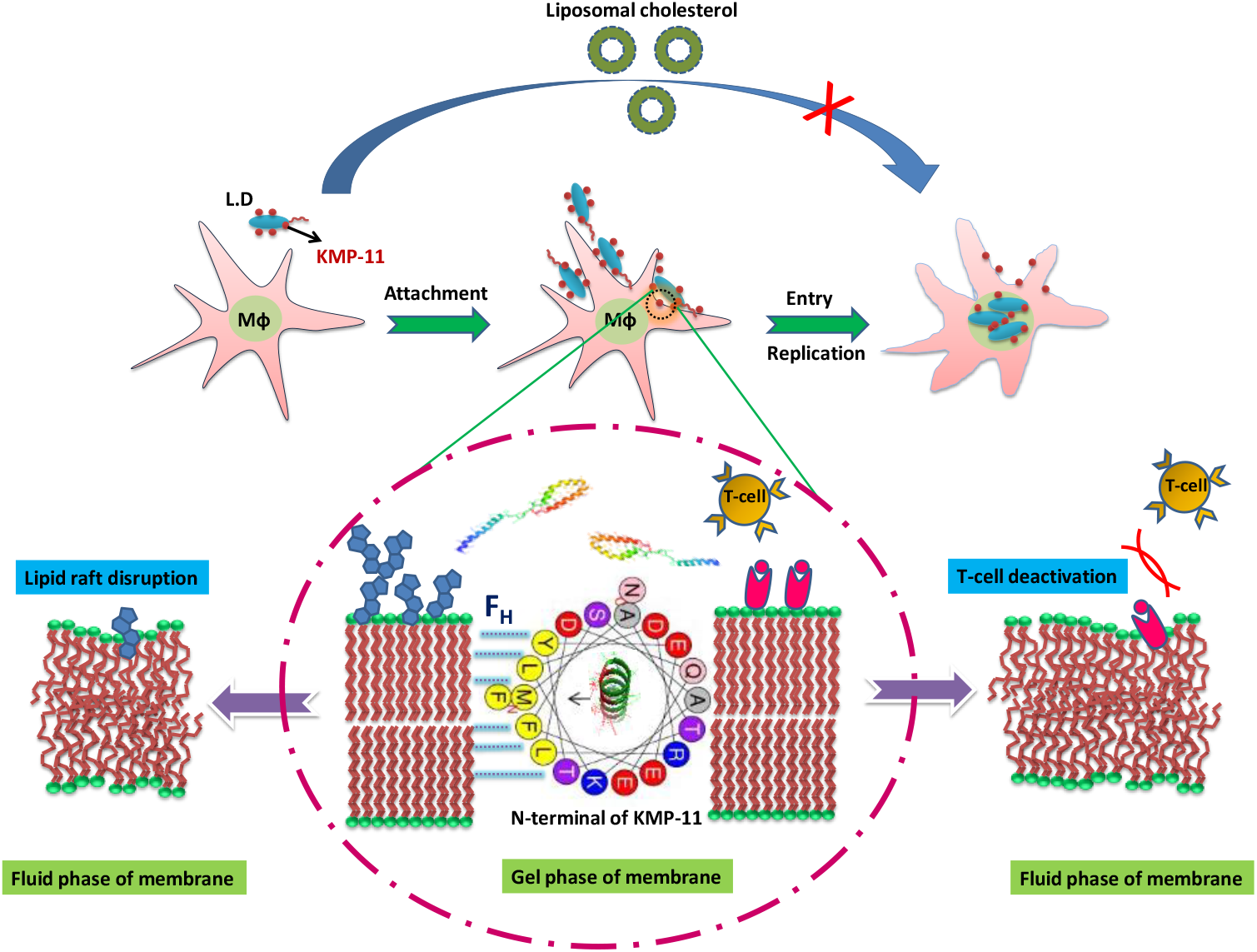
Schematic representation shows that N-terminal domain of KMP-11 binds with Mϕ membrane resulting in lipid raft disruption and T-cell deactivation. These occur through membrane deformation as a result of gel to fluid membrane phase transition. On the other hand, liposomal cholesterol transport inhibits the process of KMP-11 mediated infection pathway. Here F_H_ indicates the force generated due to the interaction of hydrophobic N-terminal face with Mϕ membrane.

The N-terminal of KMP-11 is found to play a decisive role in the membrane interaction through the mirror alignment of the polar and hydrophobic residues. The presented binding-phase change-infection model can explain results obtained in the presence of cholesterol and can be generalized by the peptide studies. We show that the binding of KMP-11 with the membrane is significantly diminished in the presence of added cholesterol. The role of cholesterol can be explained by its presumable binding as the protein is similar to the apolipoproteins. Cholesterol binding with KMP-11 competes with its membrane binding and inhibits the phase transition in membrane blocking the infection and other downstream processes. Similarly, the peptide with less μ_H_ does not bind to the membrane and does not initiate the process of LD infection.

A mathematical interpretation of the phase transition of the lipid membranes due to protein binding- comes from recent literatures, which show that the incorporation of proteins into bilayers can alter membrane tension resulting in a membrane curvature.^17–19^ Recent study using micropipette aspiration along with linear instability theory show that membrane tension and coupling between local protein density and membrane curvature mediate the shape change in the phospholipid GUV.^19^ It is likely that strong binding of KMP-11 can cause membrane instability due to protein density fluctuations. A threshold protein to lipid ratio is required for the onset of phase transition, which is clearly validated by the presented SAXS data. We find here that the gel to fluid transition occurs at or above a P/L of 0.004. It should be noted that the expressed copy number of KMP-11/ LD parasite surface is ~ 2× 10^6^. On the other hand, our calculation showed that ~ 5 ×10 ^4^ copy number of KMP-11/Mϕ (detailed calculations are provided in SI) is required for phase transition induced LD infection. Therefore, P/L, 0.004 is sufficient for Mϕ phase transition and inducing infection at physiological temperature.

Protein–lipid interactions are driven by a number of factors most notably the hydrophobic interaction. Although, it is generally accepted that globular proteins fold with a hydrophobic core and hydrophilic exterior, the centroid of the spatial distribution of amino acid residue provides the origin of moment expansion. This spatial distribution has been described by David Silverman as a two component spherical model where interacting partners had been considered as spheres.^20^ Using these considerations, we derived a “hydrophobic moment and sequence symmetry oriented phase transition model” which correlates (Equation 1) the transition temperature of the membrane (T_m_) with the number of amino acid residues present in the mirror sequence of the interacting protein stretch (N): (please refer to the SI for the details):

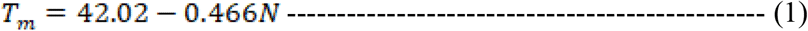

To summarize, μ_H_ of a sequence is critical to alter the conformation of the phospholipid hydrocarbon tail region through bond rotation during protein-membrane interactions leading to the change in membrane fluidity. This is expected to be a general model and should be applicable to KMP-11 and other apolipoprotein family proteins. In Table S6, we analyzed the sequence distributions of the binding domains of different apolipoproteins whose structures are known. We found a strong connection between the N values (no. of residues constituted the mirror stretch) and the apparent binding free energy of the proteins, an observation which validates the present model. It is to be noted that multiple sequence alignment of KMP-11 showed sequence similarity with different known disease causing membrane interacting proteins (Figure S14-S35). It was found that N-terminal mirror domain of KMP-11 remains conserved in all the disease protein sequences (Figure 6a). In addition, we show the disease promoting stretches of different membrane binding proteins in Figure S8. From the analyses of the membrane interacting domains, it is clear that all these stretches contain appreciable N values. It can thus be concluded that there would be a threshold N value (probably N= 5) which is necessary for the disease promoting proteins to enhance the infection propensity through membrane destabilizing strategy (Figure 6b). Interestingly, the interacting domain of KMP-11 remains preserved in other pore forming and disease promoting proteins and most of the proteins contain this sequence stretch in their pore forming regions (Figure 6a and Figure S14).

**Figure 6:**
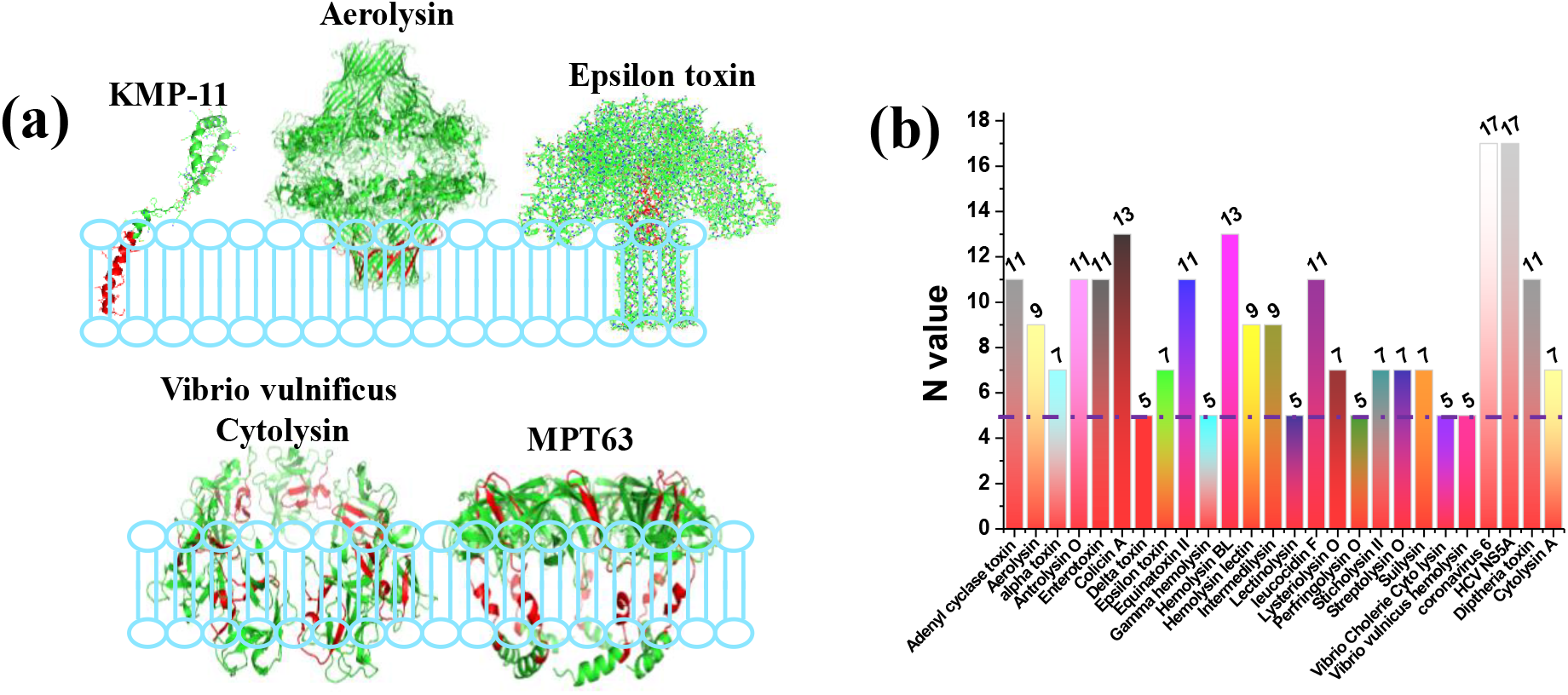
The sequence similarity between KMP-11-membrane interacting region with the membrane interacting stretches of other disease promoting proteins. **(a)** It was found that N-terminal region of KMP-11 remained conserved in all the other proteins (these conserved sequences were marked red).(b) Plot of N-values with respect to different protein species which are responsible for membrane binding and disease promotion.

Collectively, we demonstrated here that a novel mechanism of protein binding induced membrane phase transition facilitates LD infection. We also develop general framework of this mechanism by suggesting that significant N-value of the membrane interacting domain would be globally essential for the disease causing proteins which facilitate the disease process through membrane destabilization strategy. This study will also be helpful towards the development of therapeutic interventions by means of targeting this particular disease promoting sequence.

## Materials and Methods

### Materials

Dipalmitoylphosphatidyl choline (DPPC) and Dehydroergosterol were purchased from Avanti Polar Lipids Inc. (Alabaster, AL, USA). DiIC-18, NBD-C12 PC, Laurdan dye were obtained from Invitrogen (Eugene, Oregon, USA). All other necessary chemicals were obtained from Aldrich (St. Louis, USA) and Merck (Mumbai, India).

### Measurement of fluorescence anisotropy (FA)

Membrane fluidity was measured using the method of Shinitzky and Inbar.^21–22^ Briefly, the fluorescence probe DPH was dissolved in tetrahydrofuran at 2 mM concentration. To 10 ml of rapidly stirring PBS (pH 7.2), 2 mM DPH solution was added. For labeling, 10^6^ cells were mixed with an equal volume of DPH in PBS (C*f* 1 μM) and incubated for 2 h at 37°C. Thereafter the cells were washed thrice and resuspended in PBS. The DPH probe bound to the membrane of the cell was excited at 365 nm and the intensity of emission was recorded at 430 nm using a spectro-fluorometer. The FA value was calculated using the equation:

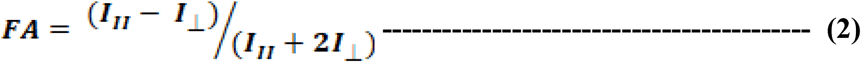

Where, I_II_ and *I*_⊥_ are the fluorescence intensities oriented, respectively, parallel and perpendicular to the direction of polarization of the excited light.

### Measurement of IL-2

RAW264.7 cells were culture overnight in a 35mm tissue culture plate (1×10^6^ cells/ml). On the next day the cells were co-cultured with LMR7.5 T cell hybridoma specific for *Leishmania* antigen for 48 hr in presence of 10μg/ml LACK antigen along with KMP11 (low dose 5μM and high dose 50 μM) respectively. After incubation cell free supernatant was collected and subjected to sandwich ELISA using IL-2 specific antibody (BD biosciences, CA, USA) as per manufacturer instruction.

### Parasite Count Measurements

RAW264.7 cells were cultured (10,000 cells/ well) in chamber slide with 200 μl/ well RPI1640 media supplemented with 10% FCS overnight. After overnight incubation, cells were treated with KMP11 and subsequently infected with *Leishmania donovani* promastigote (MHOM/IN/1983/AG83) (1:10 macrophage to parasite ratio). After 24 h, cells were washed and stained with Giemsa staining solution and the number of intracellular amastigotes were calculated and represented per 100 macrophages.

### Confocal Microscopy to detect membrane lipid raft

RAW264.7 cells were cultured (10,000 cells) on coverslip in RPI1640 media supplemented with 10% FCS overnight. After incubation cells were treated with KMP11 and subsequently infected with *L. donovani* parasite for 4 hr and washed with 1xPBS. The cells were then processed with lipid raft markers using Vybrant Alexa Fluor 488 Lipid Raft Labelling Kit (ThermoFisher Scientific, Massachusetts, USA) following manufacturers instruction and then mounted in vectashield mounting media with DAPI and subjected to observation under con-focal microscope.

### Liposomal-cholesterol preparation and delivery

Liposomes were prepared with cholesterol and phosphatidylethanolamine at a molar ratio of 1.5:1. A thin dry film of lipids (5.8 mg cholesterol and 8.0 mg phosphatidylethanolamine) was dispersed in 1 ml of RPMI 1640 and sonicated at 4°C three times, 1 min each, at maximum output. To alter the fluidity of cells, 10^5^ intact cells were incubated with liposomes for 12 h at 37°C. The cells with altered fluidity were then washed three times in serum-free RPMI 1640 medium and finally resuspended in 10% FCS containing RPMI 1640.

### Preparation of small unilamellar vesicles (SUVs) from DPPC

An appropriate amount of DPPC lipid in chloroform (concentration of stock solution is 25 mg mL^−1^) was transferred to a 10 ml glass bottle. The organic solvent was removed by gently passing dry nitrogen gas. The sample was then placed in a desiccator connected to a vacuum pump for a couple of hours to remove traces of the leftover solvent. A required volume of 20 mM sodium phosphate buffer at pH 7.4 was added to the dried lipid film so that the final desired concentration (10 mM) was obtained. The lipid film with the buffer was kept overnight at 4°C to ensure efficient hydration of the phospholipid heads. Vortexing of the hydrated lipid film for about 30 min produced multilamellar vesicles (MLVs). Long vortexing was occasionally required to make uniform lipid mixtures. This MLV was used for optical clearance assay. For preparing the small unilamellar vesicles, MLVs was sonicated using a probe sonicator at an amplitude 45% for 30 minutes and after that sample was centrifuged at 5000 rpm to sediment the tungsten artifacts and finally it was filtered by 0.22μm filter unit. Size of the small unilamellar vesicles was measured by DLS and the average diameter was found to be ~ 70 nm.

### Site directed mutagenesis

Wild type KMP-11 gene cloned previously in pCMV-LIC vector was further sub-cloned in pET28a vector between NcoI and BamHI using the following set of primers: Fwd: 5’-CCCCATGGCCACCACGTACGAGGAG-3’ and Rev: 5’-AAGGATCCCTCCTGATGATGATGATGATGCTTGGAACGGGTACTGCGCAGC-3’. Site directed mutagenesis was performed with the help of Quick-change Site-Directed Mutagenesis kit (Stratagene, Agilent Technologies) to generate several single tryptophan (W) mutants, using the wild type KMP-11 construct as a template. All the mutations were confirmed by DNA sequencing.

### Purification of the WT and Mutants of KMP-11

Recombinant KMP-11 constructs (both WT and mutants) were expressed and purified using Ni-NTA affinity chromatography. For this purpose, a Qiagen-supplied protocol (Qiaexpressionisttm; Qiagen, Germany) was used after slight modifications. The modifications included the use of 20 mM imidazole in both lysis and wash buffers during cell lysis. The purified protein fractions were checked using 15% sodium dodecyl sulfate-polyacrylamide gel electrophoresis gel electrophoresis. The collected fractions containing the protein were dialyzed using 20 mM sodium phosphate buffer at pH 7.4 to remove excess imidazole. The concentration of the protein was determined using the BCA Protein Assay Kit (Pierce, ThermoScientific).

### Structure Modeling of KMP-11

The structure modeling of KMP-11 has been discussed earlier.^23^ Sequence analysis of KMP-11 using NCBI protein−protein BLAST does not show any close homolog with the available solved structures. However, a profile-based search performed using PSI BLAST indicated the presence of few remote homologs, whose crystal structures are available. As the sequence identities of the remote homologs are low, we have used a composite approach using iterative threading assembly refinement (ITASSER) of the Zhanglab server, which combines various techniques, such as threading, ab-initio modeling, and atomic level structure refinement methods. From the I-TASSER analyses, we have chosen a model structure that provides the best confidence score.

### Binding Assay Using Fluorescence Spectroscopy

We have used a fluorometric assay for studying the binding of WTKMP-11 and its tryptophan mutants with SUVs composed of DPPC. All samples were prepared in 20 mM sodium phosphate buffer at pH 7.4. A set of samples were prepared using a 1mM concentration of uniformly synthesized lipid vesicles. In each sample vial, 0.5 wt% of membrane-specific DiI C-18 dye was added, and the samples were kept at 37 °C for overnight incubation. Subsequently, the required amount of protein was added into the vials by maintaining the lipid/protein molar ratio between 1:0 and 50:1. The samples were then incubated at room temperature (25 °C) for 2 h. The steady-state fluorescence emission spectra of the dye were recorded at an excitation wavelength of 600 nm. The peak intensity values at 683 nm for DPPC were plotted with protein concentration. The data have been fitted using the sigmoidal Hill equation, as follows:

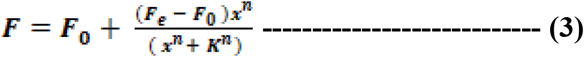

Where, *F* and *F*_*0*_ refer to the fluorescence intensities of DiI C-18 in the presence and absence of protein, respectively. *F*_*e*_ denotes the minimum intensity in the presence of a higher concentration of protein, and *K* is the equilibrium dissociation coefficient of the lipid−protein complex. *n* is the Hill coefficient, which measures the cooperativity of binding, and *x* is the concentration of the protein. PTI fluorimeter (Photon Technology International, Inc) and a cuvette with a 1 cm path length were used for the fluorescence measurements.

### Optical clearance assay

WT KMP-11 solution [10 mM NaPi, 100 mM NaCl, and 1 mM EDTA (pH 7.4)] was incubated in a quartz cuvette in an HP8453 spectrophotometer (Hewlett-Packard) at 37°C for 10 min. DMPC MLVs kept at RT were added (5% of total solution volume) to the cuvette and quickly pipetted several times to mix. The absorbance at 500 nm was recorded for 20 minutes using an HP845x UV−visible ChemStation in kinetics mode (NHLBI Biophysics Core).Typical protein /lipid MLV ratio was kept 1:200 for measurement.

### Circular Dichroism Measurement

Near and Far-UV CD spectra of WT KMP-11 in absence and presence of DPPC SUVs were recorded using a JASCO J720 spectro-polarimeter (Japan Spectroscopic Ltd.). Far-UV CD measurements (between 200 and 250 nm) were performed using a cuvette of 1 mm path length. A protein concentration of 10μM was used for CD measurements. The scan speed was 50 nm min^−1^, with a response time of 2 s. The bandwidth was set at 1 nm. Three CD spectra were recorded in the continuous mode and averaged. Typical protein-lipid ratio was maintained 1:200.

### Phase state analysis by steady state and time resolved fluorescence

Phase state of DPPC bilayer was monitored by using PTI fluorimeter for the measurement of steady state fluorescence at different temperatures in absence and presence of KMP-11 protein ratio-metric study was also done. Red Edge Excitation shift of NBD-labelled DPPC in absence and presence of protein was also determined using steady state fluorescence by applying different excitation wavelengths (465,475,485,495,505 & 515 nm).

Trans-bilayer movement of symmetrically labelled NBD-DPPC bilayer was measured from the fluorescence quenching by using dithionite as reducing agent of NBD fluorophore. This fluorescence decline corresponds to the dithionite-mediated reduction of analogs localized in the outer membrane leaflet. Subsequently, fluorescence intensity adopted a new plateau representing the analogs on the inner leaflet. The very slow decrease of fluorescence indicates that the dithionite permeation across the membrane and/or the analog flip-flop were negligible under these conditions within the time scale of the experiment. From the fluorescence intensities before (F_0_) and after (F_R_) addition of dithionite, the relative fraction of the analog on the outer leaflet (P_o_) was estimated using the following equation:

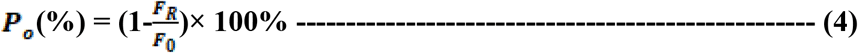

Upon addition of Triton X-100 all analogs became accessible to dithionite, resulting in a complete loss of fluorescence.

Fluorescence lifetimes were measured using time-correlated single-photon-counting (TCSPC) equipment (Fluoro cube Fluorescence Lifetime System by Horiba JovinYvon, Japan) using a picosecond pulsed diode laser with excitation at 488nm for NBD-DPPC. The total intensity decay curves, I(t), were fitted to tri exponential for the NBD-DPPC systems.

### SAXS experiment

Samples for small-angle X-ray scattering (SAXS) studies were taken in 1.0 mm glass capillaries (Hampton Research) and were flame sealed. SAXS data were collected using a HECUS S3-Micro system, equipped with a 1D position sensitive detector. Exposure times varied from 60 to 90 minutes. Data were collected over a range of the magnitude of the scattering vector (q) from 0.17 to 3.7 nm ^−1^. Error bar in the measurement of the lamellar periodicity (d) varied from 1.70 nm at very low q to 0.02 nm at very high q.

### FTIR spectroscopy

FTIR experiment was performed for determination the morphological changes of DPPC bilayer in presence and absence of KMP-11 using Bruker Tensor 27 FTIR spectrometer in liquid mode. For FTIR measurements, we considered the frequency range from 1280cm^−1^ and 1440 cm^−1^ which corresponds to the methylene wagging band (CH_2_) of DPPC and spectral position at 1367 cm^−1^ indicated the kink and gauche rotameric conformational signature of lipid. Using Gaussian fit, we measured the percentages of different conformers in absence and presence of KMP-11 and its other tryptophan mutants.

### Tryptophan quenching experiment

Steady state fluorescence spectroscopy and acrylamide quenching measurements in free and in membrane bound conditions were carried out using a PTI fluorimeter (Photon Technology International, USA). A cuvette with 1cm path length was used for the fluorescence measurements. For the tryptophan fluorescence quenching experiments, an excitation wavelength of 295 nm was used to eliminate the contributions from the tyrosine fluorescence. Fluorescence data were recorded using a step size of 1 nm and an integration time of 1sec.Excitation and emission slits were kept at 5nm in each case. Emission spectra between 305nm and 450 nm were recorded in triplicate for each experiment. Typical protein concentration of 10 μM was used for each quenching experiment and 1:200 protein-lipid molar ratio was maintained. The protein solutions were incubated at room temperature for 1 hour and then titrated using a stock of 10M acrylamide. Necessary background corrections were made for each experiment.

### Acrylamide Quenching Data Analysis

Assuming I & Io represent tryptophan fluorescence intensity of the proteins in the presence and absence of acrylamide concentration [Q], Stern-Volmer Equation^23^ can be represented as follows:

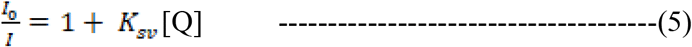

*K*_*sv*_ is the Stern-Volmer constant, which can be determined from the slope of the linear plot of I_0_/I vs. acrylamide concentrations [Q].

### Energy transfer measurement

All experiments were done using small unilamellar vesicles (SUVs) containing 1280 nmol of DPPC with required amount of DHE (4 mol %). Tryptophan fluorescence of the four mutants (Y5W, Y48W, F62W and Y89W) was monitored using steady state fluorescence technique. The tryptophan residue of the mutant served as the donor whereas DHE was used as the acceptor. The energy transfer efficiencies were calculated using the equation^24^ (Lakowicz, 1999):

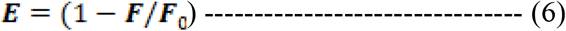

Where E is the efficiency of energy transfer and F and F_o_ are fluorescence intensities of the donor (tryptophan of KMP-11 mutants) in the presence and absence of the acceptor (DHE), respectively. Using FRET, the distance “r” between tryptophan residues of different tryptophan residues and membrane bound DHE could be calculated by the equation.

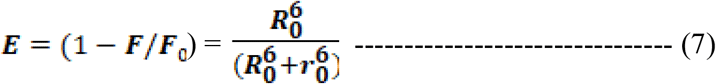

R_0_ (Forster radius), measured in Å unit, is the critical distance when the efficiency of transfer is 50%; r_0_ is the distance between donor and acceptor.

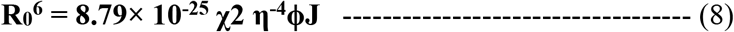

***χ***^**2**^ is the orientation factor related to the geometry of the donor and acceptor of dipoles and 2 /3 for random orientation as in fluid solution; n is the average refractive index of medium in the wavelength range where spectral overlap is significant; ϕ is the fluorescence quantum yield of the donor; J is the effect of the spectral overlap between the emission spectrum of the donor and the absorption spectrum of the acceptor, which can be calculated by the equation:

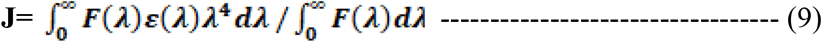

where, F(λ) is the corrected fluorescence intensity of the donor in the wavelength range from λ to λ + Δλ; ε(λ) is the extinction coefficient of the acceptor at λ.

### Peptide synthesis

Our three designed peptides (**MATTYEEFSAKLDRLDQEF, MATTYEEFSAKLDLRDQEF and MATTYEEFSALKLDRFDQE)** containing different hydrophobic moments were synthesized and supplied by Abgenex (KRIC distributor). Peptides are of good quality and ~ 95% pure.

## Supporting Information

Mathematical exposition of ‘***Hydrophobic Moment and sequence symmetry oriented phase transition model’,*** and the calculation of the number of KMP-11 molecules needed for Mϕ phase transition have been described. Additional Figures S1-S34 and Table S1-S8 have also been provided.

## Acknowledgement

Author A.Sannigrahi acknowledges University Grant Commission, Govt. of India for providing senior research fellowship. Author S.Roy acknowledges J C Bose Fellowship (Number: SB/S2/JCB-65/2014). S. Karmakar acknowledges the financial support from DBT-funded research project (BT/PR8475/BRB/10/1248/2013). K Chattopadhyay acknowledges funding from the CSIR network project grant HOPE. We thank Professors Carl Frieden and Elliot Elson of Washington University School of Medicine for their critical comments and suggestions on manuscript. We thank Professor Roop Mallik of the Tata Institute of Fundamental Research, Mumbai for multiple exciting discussions on this study, which were essential for the shaping up of this manuscript. We thank the director, CSIR-IICB for help and encouragements.

## References

1. Pearson, R. D.; Wheeler, D. A.; Harrison, L. H.; Kay, H. D., The immunobiology of leishmaniasis. Reviews of infectious diseases 1983, 5 (5), 907–927.

2. Jardim, A.; Hanson, S.; Ullman, B.; McCubbin, W.; Kay, C.; Olafson, R., Cloning and structure-function analysis of the Leishmania donovani kinetoplastid membrane protein-11. Biochemical journal 1995, 305 (1), 315–320.

3. Fuertes, M. A.; Berberich, C.; Lozano, R. M.; Gimenez- Gallego, G.; Alonso, C., Folding stability of the kinetoplastid membrane protein- 11 (KMP- 11) from Leishmania infantum. European journal of biochemistry 1999, 260 (2), 559–567.

4. Jardim, A.; Funk, V.; Caprioli, R.; Olafson, R., Isolation and structural characterization of the Leishmania donovani kinetoplastid membrane protein-11, a major immunoreactive membrane glycoprotein. Biochemical Journal 1995, 305 (1), 307–313.

5. Mukhopadhyay, S.; Sen, P.; Majumder, H. K.; Roy, S., Reduced expression of lipophosphoglycan (LPG) and kinetoplastid membrane protein (KMP)-11 in Leishmania donovani promastigotes in axenic culture. The Journal of parasitology 1998, 644–647.

6. Stebeck, C. E.; Baron, G. S.; Beecroft, R. P.; Pearson, T. W., Molecular characterization of the kinetoplastid membrane protein-11 from African trypanosomes. Molecular and biochemical parasitology 1996, 81 (1), 81–88.

7. Gwynne, J.; Brewer, H.; Edelhoch, H., The molecular behavior of apoA-I in human high density lipoproteins. Journal of Biological Chemistry 1975, 250 (6), 2269–2274.

8. Ghosh, J.; Guha, R.; Das, S.; Roy, S., Liposomal cholesterol delivery activates the macrophage innate immune arm to facilitate intracellular Leishmania donovani killing. Infection and immunity 2014, 82 (2), 607–617.

9. Sannigrahi, A.; Nandi, I.; Chall, S.; Jawed, J. J.; Halder, A.; Majumdar, S.; Karmakar, S.; Chattopadhyay, K., Conformational switch driven membrane pore formation by Mycobacterium secretory protein MPT63 induces macrophage cell death. ACS chemical biology 2019.

10. Learmonth, R. P.; Gratton, E., Assessment of membrane fluidity in individual yeast cells by laurdan generalised polarisation and multi-photon scanning fluorescence microscopy. In Fluorescence Spectroscopy, Imaging and Probes, Springer: 2002; pp 241–252.

11. Chakraborty, D.; Banerjee, S.; Sen, A.; Banerjee, K. K.; Das, P.; Roy, S., Leishmania donovani affects antigen presentation of macrophage by disrupting lipid rafts. The Journal of Immunology 2005, 175 (5), 3214–3224.

12. Schroit, A.; Gallily, R., Macrophage fatty acid composition and phagocytosis: effect of unsaturation on cellular phagocytic activity. Immunology 1979, 36 (2), 199.

13. A. Sannigrahi, P. M., S. Karmakar and K. Chattopadhyay, Interaction of KMP-11 with Phospholipid Membranes and Its Implications in Leishmaniasis: Effects of Single Tryptophan, Mutation and Cholesterol. J. Phys. Chem B 2017, 121, 1824–1834.

14. Stevenson, P.; Tokmakoff, A., Time-resolved measurements of an ion channel conformational change driven by a membrane phase transition. Proceedings of the National Academy of Sciences 2017, 114 (41), 10840–10845.

15. Lewis, R. N.; McElhaney, R. N., Membrane lipid phase transitions and phase organization studied by Fourier transform infrared spectroscopy. Biochimica et Biophysica Acta (BBA)-Biomembranes 2013, 1828 (10), 2347–2358.

16. Raghuraman, H.; Shrivastava, S.; Chattopadhyay, A., Monitoring the looping up of acyl chain labeled NBD lipids in membranes as a function of membrane phase state. Biochimica et Biophysica Acta (BBA)-Biomembranes 2007, 1768 (5), 1258–1267.

17. Elliot L. Elson, E. F., John E. Dolbow, Guy M. Genin, Phase separation in biological membranes: integration of theory. Biophys. J 2010, 39, 207–226.

18. Mouritsen, O. G., MATTRESS MODEL OF LIPID-PROTEIN INTERACTIONS IN MEMBRANES. Biophys. J 1984, 46, 141–153.

19. Baumgart, Z. S. a. T., Membrane tension and peripheral protein density mediate membrane shape transitions. Nature communication 2015, 5974 (6), 1–8.

20. Silverman, B. D., Hydrophobic moments of protein structures: Spatially profiling the distribution. Proceedings of the National Academy of Sciences 2001, 98 (9), 4996–5001.

21. Inbar, M., Fluidity of membrane lipids: a single cell analysis of mouse normal lymphocytes and malignant lymphoma cells. FEBS letters 1976, 67 (2), 180–185.

22. Shinitzky, M.; Henkart, P., Fluidity of cell membranes—current concepts and trends. In International review of cytology, Elsevier: 1979; Vol. 60, pp 121–147.

23. Lakowicz, J., Principles of fluorescence microscopy. Kluwer Academic, New York: 1999.

24. Lakowicz, J., Principles of fluorescence spectroscopy. Kluwer Academic, New York. Principles of fluorescence spectroscopy. 2nd ed. Kluwer Academic, New York. 1999, -.

