## Supporting Information for "Protein induced membrane phase transition facilitates leishmania infection"

### Table of contents

*1. Mathematical Exposition of phase transition model* **S 03-S 07**

*2.Calculation of KMP-11 for phase transition* **S 08-S 09**

*3. Tables S1-S8 describing Fluorescence lifetimes,binding assays,*

*Quenching measurements,REES and laurdan data, energy transfer,*

*KMP-11 sequence stretches, binding energies, binding*

*of apolipoprotein family proteins, sequences of disease*

*causing proteins and their mirror sequence N values*

*and the primers design* **S 10-S 17**

*5. Laurdan based fluidity measurement in presence and absence of KMP-11* **S18-S18**

*6. Membrane binding and conformation study of KMP-11* **S 19-S 19**

*7. DPPC laurdan data at different P/L and effect of cholesterol* **S 20-S 21**

*8. Optical clearance study and REES study* **S 22-S 23**

*9. FTIR of membrane and positioning of KMP-11 on membrane* **S 24-S25**

*10. FRET study* **S26-S26**

*11. Peptide membrane interactions* **S27-S27**

*12. Sequence similarity analysis* **S29-S49**

*13. References* **S50-S50**

Mathematical exposition of '***Hydrophobic moment and sequence symmetry oriented phase transition model***':

Protein–lipid interactions are driven by number of factors. Among these, hydrophobic interaction is the principal driving force. Although, it is generally accepted that globular proteins fold with a hydrophobic core and hydrophilic exterior, the centroid of the spatial distribution of amino acid residue provides the origin of moment expansion. Spatial distribution has been described by David Silverman as a two component spherical model where interacting partners had been considered as spheres.[1] Subsequently, we constructed a '***Hydrophobic moment and sequence symmetry oriented phase transition model***', in which the nature of a helical stretch was theoretically related to the phase melting temperature of phospholipids.

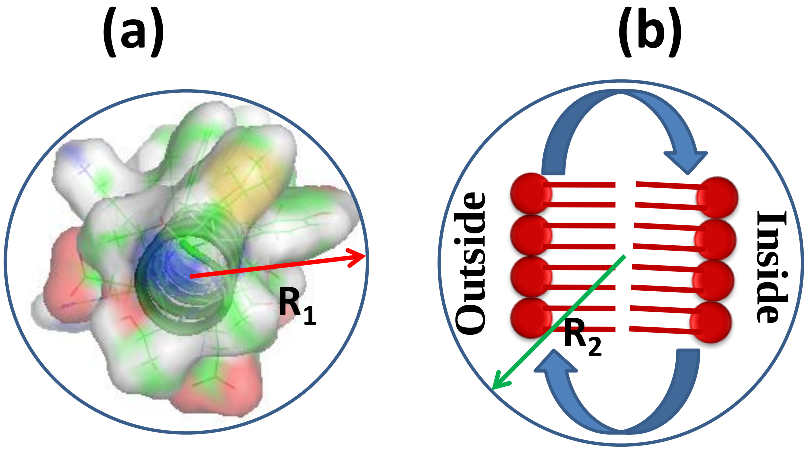


**Here, $R_1$ is variable and $R_2$is fixed.**

**Figure S1: Two interacting domains, which are considered spheres.** (a) Model Pymol structure of the N-terminal stretch of KMP-11 (1-19AA), based on which three peptides were synthesized [peptide 1, peptide2 and peptide3]. Here we assume the peptide domain as a sphere model of radius $R_1$. (b) On the other hand, considering the flip-flop dynamics of the bilayer entity, we assumed the specific interacting domain of the lipid bilayer as another sphere of radius $R_2$.$R_2$ is fixed since we are not changing any property of bilayer whereas $R_1$is variable due to the change in the sequence arrangement.

Assuming the density of hydrophobicity is ρ(R) and the radial distance from the centre of the sphere is R, it can be written as:

$\rho(R) = \alpha R^{\eta} - \beta R^{\mu}$,--------------------------(1)

Where, hydrophobic component will contribute an amount $4\pi\alpha R^{\eta} R^2 dR$ in a shell of width dR, while the contribution of the polar component is $-4\pi\beta R^{\mu} R^2 dR$.

Assuming the contribution of polar component is significantly less compared to the hydrophobic component, we can write equation (1) as

$$R= [\rho(R)/\alpha]^{1/\eta} \quad \text{------------------------(2)}$$

Jacob Israelachvili and Richard Pashley have shown that a pair interaction free energy (ΔG) abides by the following relation[2]:

$\Delta G_H = -CRDe^{-D/D_0}$--------------------(3)

Here, hydrophobic interaction is a two component system. Therefore, for a two component protein –lipid system, $R = {R_1 R_2}/{(R_1 + R_2)}$;where $R_1$(Peptide) and $R_2$(Lipid) are the radii of the interacting partners. Now equation (3) will be, $\Delta G_H = -C\left({R_1 R_2}/{R_1 + R_2}\right) De^{-\frac{D}{D_0}}$

$$= -C\left({1}/{(1/R_1 + 1/R_2)}\right) De^{-\frac{D}{D_0}} \text{----------(4)}$$

Since the lipid component is fixed and the second component (membrane interacting domain of the protein; different peptides) is variable, we can rearrange the equation (4) as

$$\Delta G_H = -CR_1 De^{-\frac{D}{D_0}}$$

$$= -CDe^{-\frac{D}{D_0}} \left(\frac{\rho(R_1)}{\alpha}\right)^{\frac{1}{\eta}}$$

$$= -CDe^{-\frac{D}{D_0}} \left(\frac{H}{\alpha A}\right)^{\frac{1}{\eta}} \text{------------------------- (5)}$$

Where, H is the net hydrophobicity and A is the area of hydrophobic face.

From our experimental and theoretical calculations, it has been established that protein-lipid interaction and phase transition is dependent on the hydrophobic moment of the proteins stretch, therefore, equation (4) can be represented as

$$\Delta G_H = -CDe^{-\frac{D}{D_0}}\left(\frac{\mu_H}{\alpha A}\right)^{\frac{1}{\eta}}\text{------------------------------------------} (6)$$

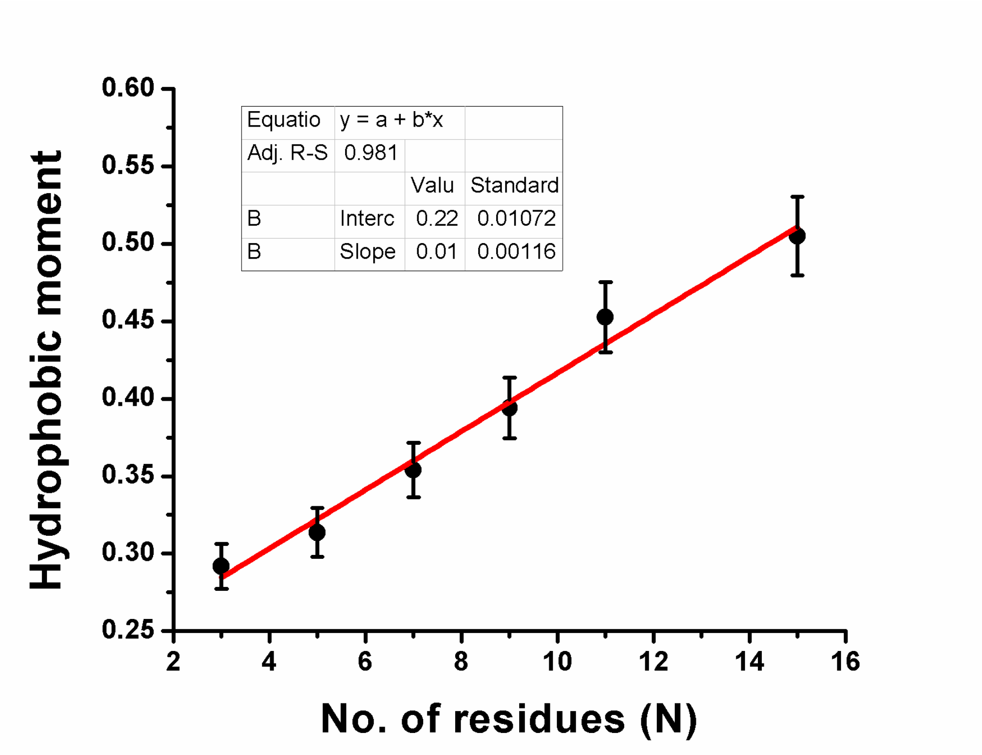


**Figure S2: Relation between hydrophobic moment and the mirror sequence constituting residue number of the membrane interacting stretch.** Linear plot of hydrophobic moment ($\mu_H$) against the number of residues constituting the mirror sequences (N) of different combinations based on polarity and hydrophobicity of the amino acid residues in the N-terminal domain of KMP-11 (Table S5).

Using the mirror sequence constituting residue number (N) and different combinations of polar (P) and hydrophobic (H) residues of KMP-11 N–terminal stretch (1-19) (Table S5), we showed the linear correlation between the hydrophobic moment, $\mu_H$ and N. The linear equation is expressed (Figure S2)as:

$$\mu_H = 0.018N + 0.227 - - - - - (7)$$

Now, combining equation (6) and (7), we get

$$\Delta G_H = -CDe^{-\frac{D}{D_0}}\left(\frac{0.018N+0.227}{\alpha A}\right)^{\frac{1}{\eta}} \text{------------------------ (8)}$$

Since hydrophobic free energy is the strongest component for the phase transition to occur, we can write, $\Delta G_H = \Delta G_T$; where $\Delta G_T$ is the free energy change during gel/fluid phase transition. Now

Thermodynamic equation of free energy would be,

$\Delta G_T = \Delta H_T - T_T \Delta S_T$ -------------------------------- (9) where $T_T$ stands for the phase transition temperature of phospholipid.

Combining equation (8) and (9), we can write,

$$T_T \Delta S_T = \Delta H_T + CDe^{-\frac{D}{D_0}}\left(\frac{0.018N+0.227}{\alpha A}\right)^{\frac{1}{\eta}}$$

Or, $T_T = \Delta H_T/\Delta S_T + \frac{CDe^{-\frac{D}{D_0}}\left(\frac{0.018N+0.227}{\alpha A}\right)^{\frac{1}{\eta}}}{\Delta S_T}$ --------------------------------- (10)

Assuming η = 1, we can write equation (10) as

$T_T = \Delta H_T/\Delta S_T + \frac{K(0.018N+0.227)}{\Delta S_T}$ ------------------------------------- (11)

Where, $K = \frac{CDe^{-\frac{D}{D_0}}}{\alpha A}$

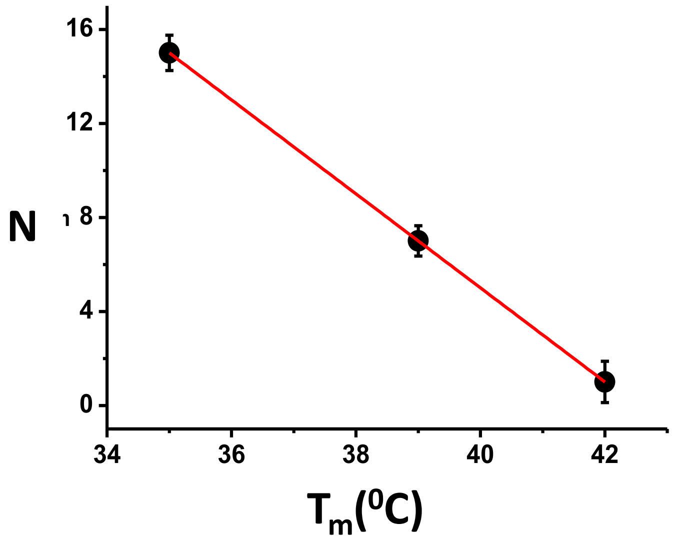


**Figure S3: Relation between mirror stretch constituting residues and chain melting temperatures of phospholipid model membranes.** Linear plot of phase transition temperature($T_m$) of DPPC against the number of residues constituting the mirror sequences (N) of different combinations based on polarity and hydrophobicity of the amino acid residues. The chain melting temperatures were obtained from the laurdan experiments of DPPC model membranes in presence of three synthetic peptides.

Linear fitting (Figure S3) of the experimental data of transition temperature ($T_m$ ) against N has given us simplified linear correlation e.g

$T_m = 42.02 - 0.466N$ ---------------------------------------------- (12)

The proposed model indicated a strong correlation between the sequence amino acid distribution and the protein mediated phase transition temperature of phospholipid membranes.

<u>**Evaluation of critical P/L ratio for phase transition of Mϕ membrane during LD infection:**</u>

It has been found that at P/L ratio> 0.004, DPPC remains as solution (Fluid phase) phase at physiological temperature ($37^0$C) which resembles the actual infection condition.

Now we want to quantify the number of KMP-11 copy number essential for inducing the phase state change of host macrophage membranes at physiological condition.

It is known that lipid head group area[3] = 63 $Å^2$

Our as prepared Vesicles (SUVs) are of radius 70 nm.

Then area of vesicles = $4\pi r^2$ = 61544 $nm^2$.

Vesicles consist of bilayer. Assuming both layer similar (layer approximation), the overall area would be= 61544 X 2= 123088 $nm^2$.

Then, no. of lipid molecules (fatty acids) in a vesicle = 123088 $nm^2$/63 $Å^2$ = 195378

Now, working concentration of DPPC lipid = 1 mM.

1 M is equivalent to 6.023 X $10^{23}$ no. of fatty acids

So, no. of vesicles in 1 mM = 6.023 X$10^{20}$/195378 =3.08 X$10^{15}$

The number of vesicles in our working solution = 3.08 X$10^{15}$

On the other hand, protein concentration = 4 μM (since, we have taken P/L = 0.004 and in this concentration phase transition occurred at below physiological temperature).

Now, 4 μM of KMP-11 is equivalent to 6.023X$10^{23}$. 4 X $10^{-6}$ = 2.4 X $10^{18}$ number of KMP-11

Therefore, in working (favorable for infection through phase transition) condition,

No. of proteins needed/ lipid vesicles = 2.4 X$10^{18}$/3.08 X$10^{15}$= 800

Considering the average size of macrophages = 5 μm which will be consisted of 1.25 X $10^{7}$ fatty acids

Now, 195378 fatty acids correspond to 800 number of KMP-11

So, 1.25 X $10^{7}$fatty acids correspond to [1.25 X $10^{7}$ / 195378].800

~ 5 X $10^{4}$ copy number of KMP-11.

According to the reports, the expressed copy number of KMP-11/ LD parasite surface is

~ 2 X $10^6$.

So, sufficient number of KMP-11 is available for the interaction and inducing infection.

**Table S1: Fluorescence lifetimes of DiI C-18 DPPC membrane in presence and absence of WT KMP-11.**

| Systems | $T_1$ | $T_2$ | $A_1$ | $A_2$ | $T_{avg}$ | $X^2$ |
|---|---|---|---|---|---|---|
| **DiI C-18-DPPC** | 3.191 | 5.154 | 0.590 | 0.410 | 4.229 | 1.091 |
| **DiI C-18-DPPC-KMP-11** | 2.412 | 4.872 | 0.630 | 0.370 | 3.746 | 1.049 |

**Table S2: Binding constants ($K_a$), cooperative indices ($n$) and Stern-Volmer constants ($K_{sv}$) of different protein mutants with DPPC model membranes.**

| Systems | $K_a$ ($M^{-1}$) | $n$ | $K_{sv}$($M^{-1}$) |
|---|---|---|---|
| **WT-DPPC** | $3.504 \times 10^6$ | 2.008 | ------ |
| **Y5W** | -------- | ------- | 4.909 |
| **Y5W-DPPC** | $5.154 \times 10^7$ | 2.117 | 2.453 |
| **Y48W** | ---------- | --------- | 1.799 |
| **Y48W-DPPC** | $8.508 \times 10^6$ | 1.945 | 1.455 |
| **F62W** | --------- | ----------- | 3.381 |
| **F62W-DPPC** | $2.630 \times 10^5$ | 1.562 | 3.177 |
| **Y89W** | ---------- | ---------- | 4.454 |
| **Y89W-DPPC** | $6.205 \times 10^6$ | 2.364 | 4.152 |

**Table S3: Chain melting temperature of DPPC membrane using using different protein (P)/Lipid(L) molar ratio as monitored by measuring the polarsiation of Laurdan and REES of NBD PC.**

| P/L ratio | Chain melting temperature using laurdan | Chain melting temperature using NBDPC |
|---|---|---|
| 0 | 44.8 | 44.4 |
| 0.0002 | 47.4 | 46.2 |
| 0.001 | 48.0 | 43.9 |
| 0.002 | 40.4 | 36.5 |
| 0.0025 | 39.5 | 37.7 |
| 0.004 | 38.7 | 33.9 |
| 0.02 | 33.5 | 33.2 |

**Table S4: Efficiency of energy transfer(E) and donor-acceptor distances ($r_0$) of different tryptophan residues of four single tryptophan mutants. Here, tryptophan was the donor and DHE was used as the acceptor molecules.**

| **Mutants** | **E** | **$r_0$ (Å)** |
|---|---|---|
| Y5W | 0.20 | 11.4 |
| Y48W | 0.06 | 14.1 |
| F62W | 0.01 | 24.7 |
| Y89W | 0.08 | 13.6 |

**Table S5: Different combinations of the amino acid residues of the N-terminal strectch of KMP-11 based on their polarity and hydrophobicity. Here, N denotes the number of amino acid residues and $\mu_H$ is the hydrophobic moments calculated by the Helical wheel model. The membrane embedded residues and the free energy changes were calculated in OPM server.**

| Serial No. | Peptide sequences | N | μH | Embedded residues | ΔG(Kcal/mole) |
|---|---|---|---|---|---|
| 1 | MATTYEEFSAKLDRLDQEF<br>HHPPHPPHPHPHPPHPPPH | 15 | 0.505 | 1,5,8,12,15,19 | -9.7 |
| 2 | MATTYEEFSAKLDRLFDQE<br>HHPPHPPHPHPHPPHHPPP | 11 | 0.453 | 1,2,5,8,12 | -7.7 |
| 3 | MATTYEEFSAKLDLRDQEF<br>HHPPHPPHPHPHPPPPHPH | 9 | 0.394 | 1,5,8,12,19 | -8.4 |
| 4 | MATTYEEFSAKLDLRDQEF<br>HHPPHPPHPHPHPHPPPPH | 7 | 0.354 | 1-5,8 | -7.3 |
| 5 | MATTYEEFSAKLLDRDFQE<br>HHPPHPPHPHPHHPPPHPP | 5 | 0.306 | 1-6,8 | -6.8 |
| 6 | MATTYEEFSAKDLRDLFQE<br>HHPPHPPHPHPPHPPHHPP | 3 | 0.292 | 1-2,4-5,8 | -6.6 |

**Table S6: Different binding domains of Apolipo-family proteins and their binding free energies.**

| Proteins | PDB ID | Binding domain sequences | N | Embedded residues | ΔG(Kcal/mole) |
|---|---|---|---|---|---|
| Apo-E4 | 1GS9 | RLLRDADDLQKRLAVYQAG<br>PHHPPHPPHPPPHHHHPHP | 7 | 164 | -3.1 |
| Apo-E | 2KC3 | MKVEQAVETEPEPELRQQTEWQSGQR<br>HPHPPHHPPPPPPPHPPPPPHPPIPP | 9 | 20 | -3.3 |
| Apo-A-I | 3r2p | AHVDALRTHLAPYSDELRQRLAAR<br>HPHPHHPPPHHPHPPPHPPPHHHP | 13 | 3-4,155-165 | -3.8 |
| Apo-C-III | 2jq3 | MQGYMKHATKTAKDALSSVQESQVA<br>HPIHHPPHHPHHPPHHPPHPPPPHH | 19 | 8-9,11-12,15-16,18-31,33,61,65 | -5.6 |
| Apo-C-I | 1IOJ | TPDVSSALDKLKEFGNTLEDKARE<br>HPPHPPHHPPHPPHIPPHPPPHPP | 19 | 1-3,14-23 | -5.8 |

**Table S7:Disease causing proteins with their membrane binding domains and respective N values.**

| DISEASE | PROTEIN | MEMBRANE BINDING STRETCH | N |
|---|---|---|---|
| Whooping cough | Adenyl cyclase toxin of B. pertusis | LFGRAPEVIARA<br>HHHHHHPHHHHH | 11 |
| Diarrhoeal diseases and deep wound infections | Aerolysin | RLFSLGQGVCGDK<br>HHHPHHPHHPHPP | 9 |
| Gas gangrene | Alpha toxin from C. perfringens | KDNSWYLAYSIPDTGES<br>PPPPHHHHHPHHPPHPP | 7 |
| Anthrax | Anthrolysin O | KVSIGGTTLYPTATISH<br>PHPHHHPPHHHPHHHPP | 11 |
| Food poisoning/gastrointestinal illnesses | Clostridium perfringens enterotoxin | QSLGDGVKDHYVDISL<br>PPHHPHHPPPHHPHPH | 11 |
| Cholera | Colicin A | EVESWVLSGIASSVAL<br>PHPPHHHPHHHPPHHH | 13 |
| Respiratory tract infection | Delta toxin | IRKEENGNVTIITQNNKQ<br>HHPPPPHPHPHHPPPPPP | 5 |
| Gas gangrene, | Epsilon toxin | TTTHTVGTSIQATAKFTVPFNETGVSLTTSYSFANTN<br>PPPPPHHPPHPHPHPHPHHHPPPHHPHPPPHPHHPPP | 7 |
| Cardiotoxic effects | Equinatoxin II | QKDRGPVATGAVGVLAYLMSD<br>PPPHHHHHPHHHHHHHHHHPP | 11 |
| Hemolytic activity | Gama hemolysin | SRTTYSDLIKRMIWPF<br>PHPPHPPHHPHHHHHH | 5 |
| endophthalmitis | Hemolysin BL | LSEIEQTNNGDTAL<br>HPPHPPPPPHPPHH | 13 |
| Hemolytic activity | Hemolysin lectin | NVLATQTLENTSSQTQE<br>PHHHPPPHPPPPPPPPP | 9 |
| Forms cytotoxic pores in CD59-positive cells, promotes adherence to and invasion of human liver cells | Intermedilysin | ATGLAWEPWRLIYS<br>HPHHHHPHHHHHHP | 9 |
| Forms cytotoxic pores in CD59-positive cells, promotes adherence to and invasion of human liver cells | Lectinolysin | EVFRSATNIG<br>PHHHPHPPHH | 5 |
| Kill leucocytes | Leucocidin F | RWNGFYWAGANY<br>HHPHHHHHHHPH | 11 |
| listeriosis | Listeriolysin O | ECTGLAWEWWRTVID<br>PPPHHHHPHHHPHHP | 7 |
| | Perfringolysin O | YDVPLTNNINVSIWGTTLYPGSST<br>HPHHHPPPHPHPHHHPPHHHHPPP | 5 |
| | Sticholysin II | SVPFDYNWYSNWWDVK<br>PHHHPHPHHPPHHPHP | 7 |
| Disrupts cytoplasmic integrity of erythrocytes, leukocytes, macrophages, platelets, epithelial cells | Streptolysin O | TGLAWEWWRKV<br>PHHHHPHHHPH | 7 |
| Cytotoxic to endothelial cells, epithelial cells, macrophages and neutrophils | Suilysin | CTGLAWEWWRTVY<br>PPHHHHPHHHPHH | 7 |
| Cholera | Vibrio cholerae cytolysin | AYKHYYVVGAHQSYH<br>HHPPHHHHHHPPPHP | 5 |
| Cholera | Vibrio vulnificus hemolysin | MAHVTLQSLSNN<br>HHPHPHPPHPPP | 5 |
| Severe Acute Respiratory Syndrome | Coronavirus Protein 6 | MFHLVDFQVTIAEILIIIIMRTFRIAIWNLDVIISSIVR<br>HHPHHHPHPHPHPPHHHHHHHHPPHPHHHHPHP | 17 |
| Hepatitis C | HCV NS5A (Hepatitis C virus) | DTSWLRDVWDWVCTVLSDFRVWLQAKLL<br>PPPHHPPHHPHHPPHHPPHPHHHPPPHH | 17 |

| | | | |
|---|---|---|---|
| Diptheria | Diptheria toxin | **SLTGTNPVFAGANYAAWAVN PHPPPPHHHHPHPHHHHHHP** | 11 |
| | Cytolysin A | **KTVEVVKNAIETADGLDLYNKYLD PPHPHHPPHHPPHPPHPHHPPHHP** | 7 |

**Table S8: Primers of the single tryptophan mutants of KMP-11**

| Tryptophan Mutations | Primers (5' → 3') |
|---|---|
| Y5W | Fwd:CCATGGCCACCACGTGGGAGGAGTTTTCGGCG Rev: GCCGAAAACTCCTCCCACGTGGTGGCCATGG |
| Y48W | Fwd: CGAGATGAAGGAGCACTGGCAGAAGTTCGAGCGCAT Rev: ATGCGCTCGAACTTCTCCCAGTGCTCCTTCATCTCG |
| F62W | Fwd:GCTCGTGCATCTTCTTGTTCCACTTCTCTGTGTGTTCCTT Rev:AAGGAACACACAGAGAAGTGGAACAAGAAGATGCACGAGC |
| Y89W | Fwd: AGCAGAAGGCTGCGCAGTGGCCGTCCAAGC Rev: GCTTGGACGGCCACTGCGCAGCCTTCTGCT |

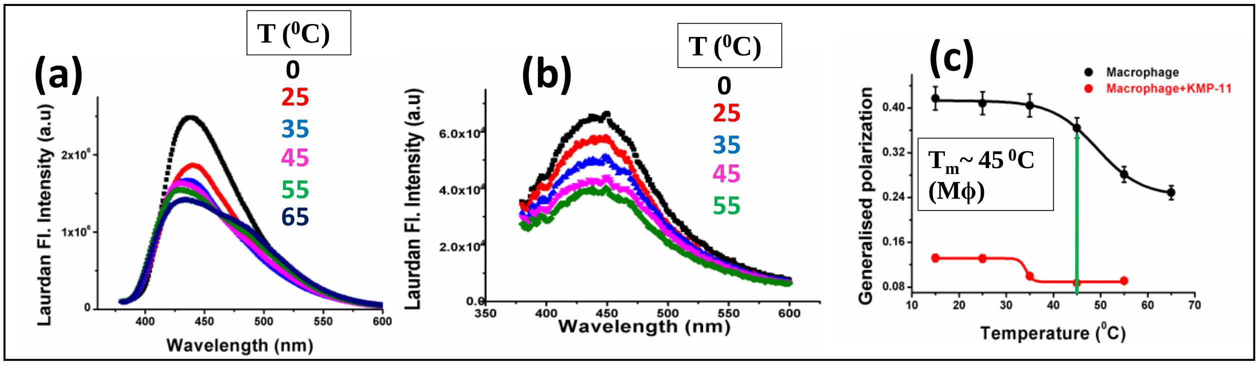


**Figure S4: Measurement of Mϕ membrane fluidity in absence and presence of KMP-11.**Plot of laurdan fluorescence intesity of labeled Mϕ with increasing temperature in the absence (a) and presence (b) of KMP-11. (c) Generalised polarization (GP) of laurdan labeled macrophage membrane with respect to temperature in absence (black) and presence (red) of KMP-11.Typically, concentration of KMP-11 was taken 50 μM.

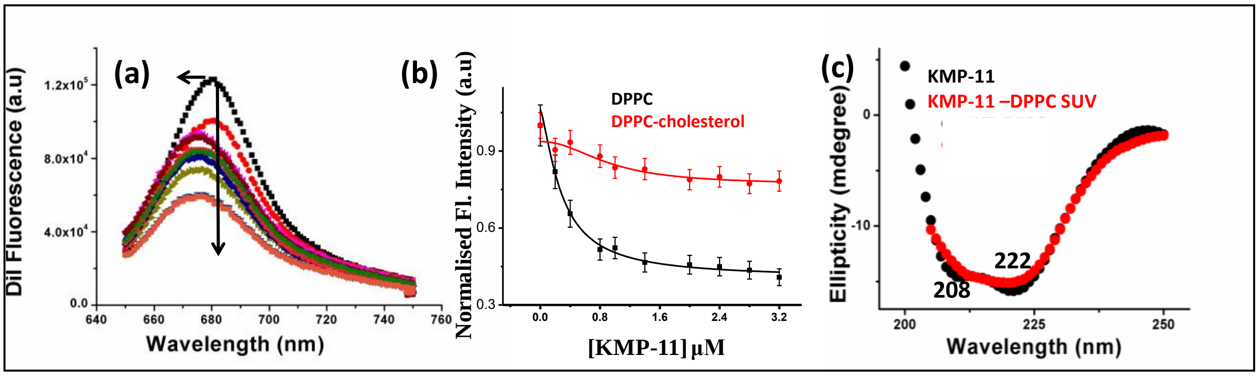


**Figure S5: KMP-11-model membrane binding and structural characteristics.** (a) DiI fluorescence intensity Vs wavelength spectra with increasing concentrations of proteins.(b)Binding of KMP-11 with DPPC model membranes in the absence (black) and presence (red) of cholesterol was monitored by decrease in DiI C-18fluorescence intensity. (c)Far UV-CD spectra of KMP-11 in the absence and presence of DPPC membrane.

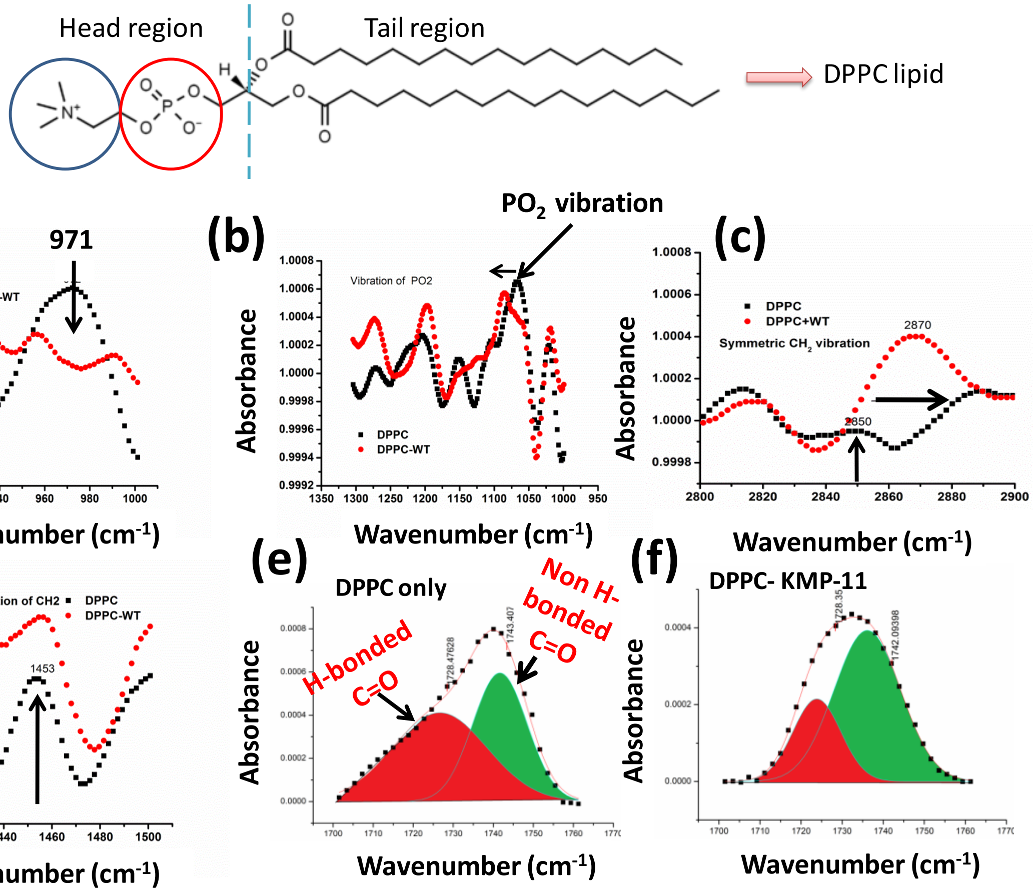


**Figure S6: Membrane perturbations study due to KMP-11-lipid binding.**FTIR signatures of (a) C-N-C,(b) $PO_2$, (c) symmetric vibrations of $CH_2$, (d) symmetric deformation of $CH_2$both in absence and presence of KMP-11. Carbonyl (C=O) stretching vibrations in absence (e) and presence of (f) KMP-11.

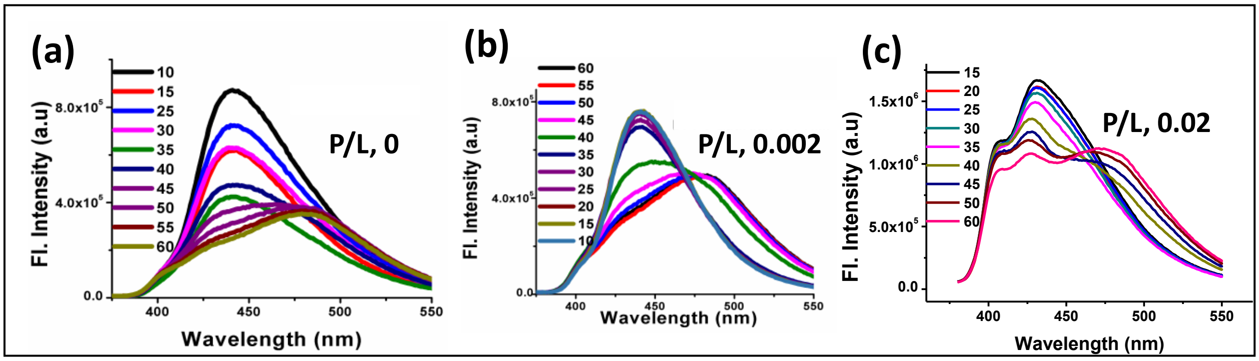


**Figure S7:Model membrane fluidity measurements in presence of KMP-11.**DPPC laurdan fluorescence in absence and presence of KMP-11 with increasing temperature when P/L ratios were kept at (a) 0, (b) 0.002 and (c) 0.02.

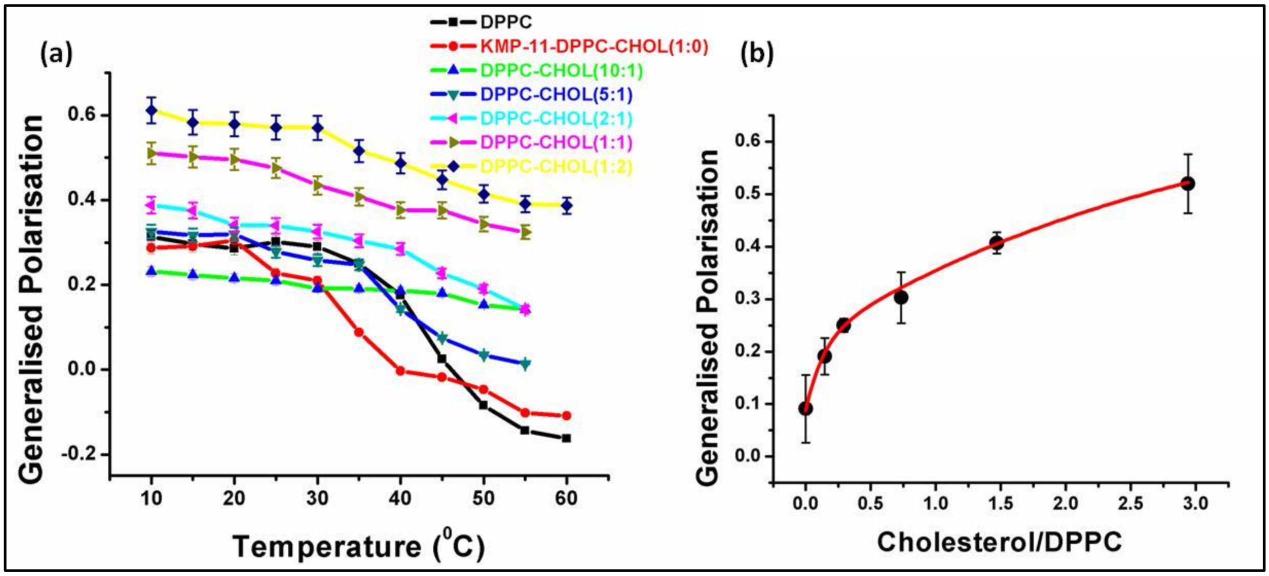


**Figure S8: Modulation of KMP-11 driven membrane fluidity in presence of different mole percentage of cholesterol.** (a) Laurdan generalized polarization of model DPPC SUVs containing different molar percentage of cholesterol with respect to temperature. All the samples of DPPC-cholesterol SUVs were treated with WT KMP-11 at Lipid/protein ratio (50:1). (b) Plot of Laurdan generalized polarization of DPPC SUVs against cholesterol/DPPC ratio. These polarization values were taken at $37^0$C temperature under the treatment of WT KMP-11 (Lipid/protein ratio was 50:1).

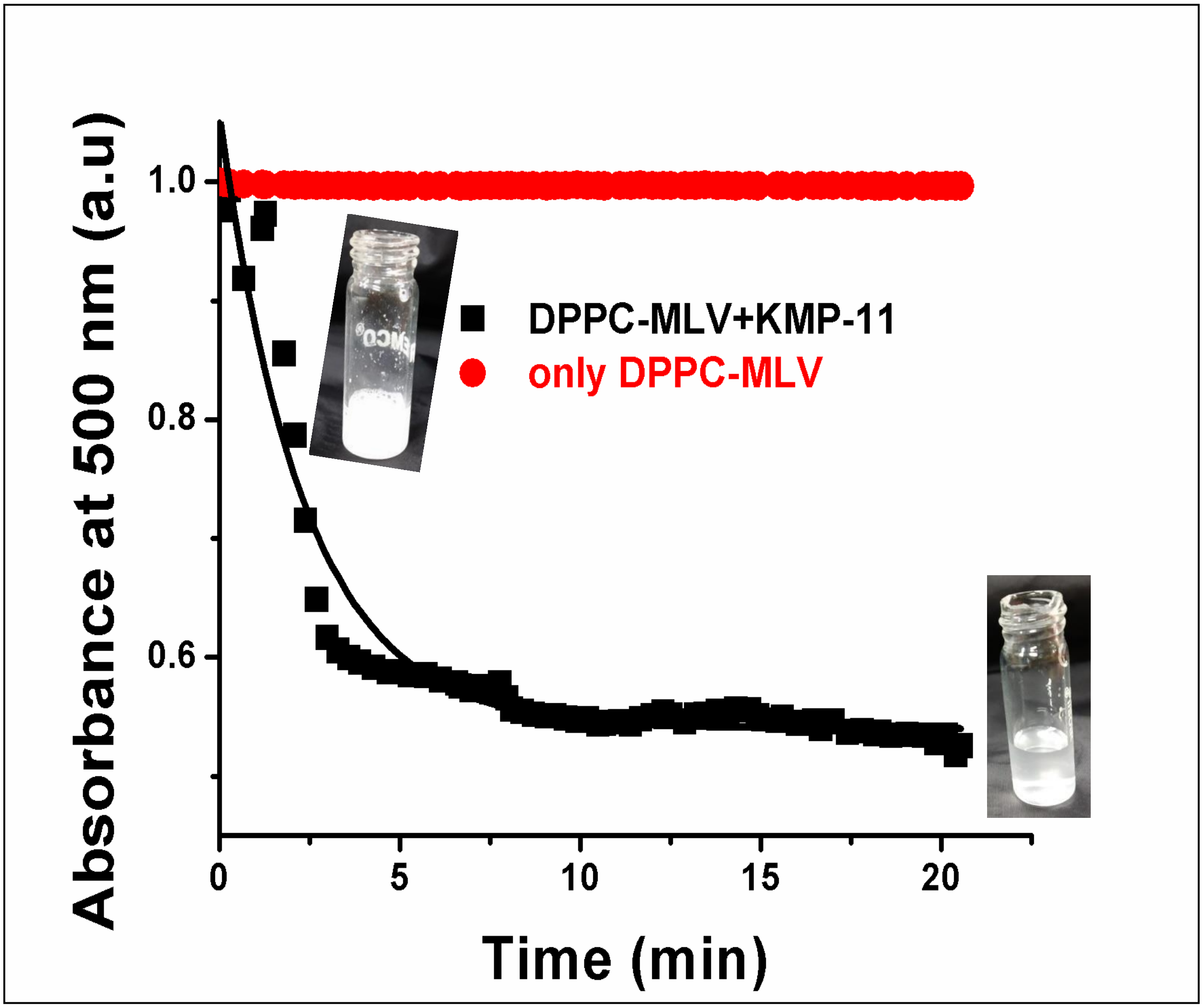


Figure S9:**Optical clearance assay of DPPC MLV in absence and presence of KMP-11 with time**. Absorbance of DPPC MLV was recorded at 500 nm with and without KMP-11 at 37 $^0$C.

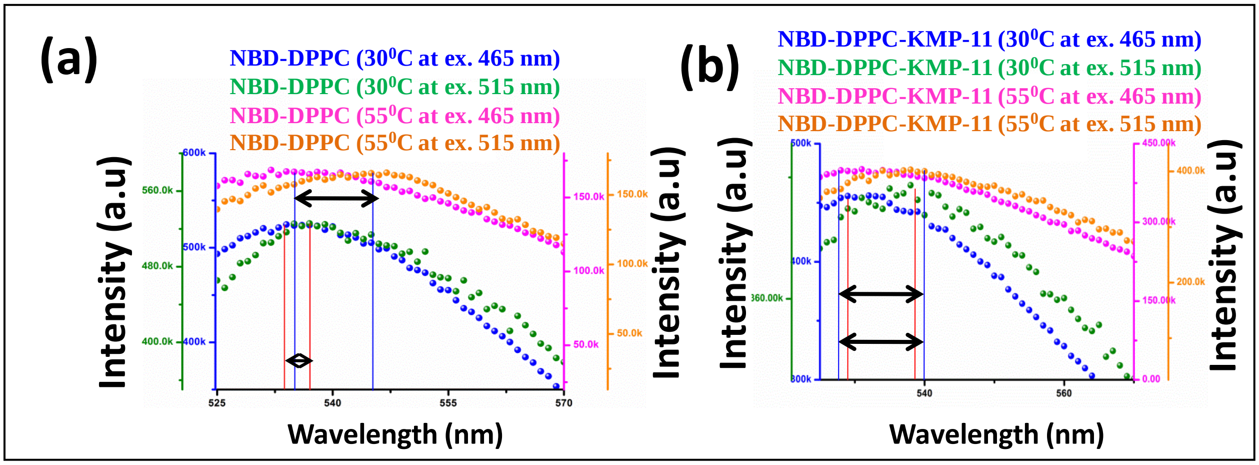


Figure S10:Maximum wavelength peak shift at temperature 30$^0$C and 55 $^0$C to monitor the REES of NBD-DPPC in absence (a) and presence (b)of WT KMP-11.

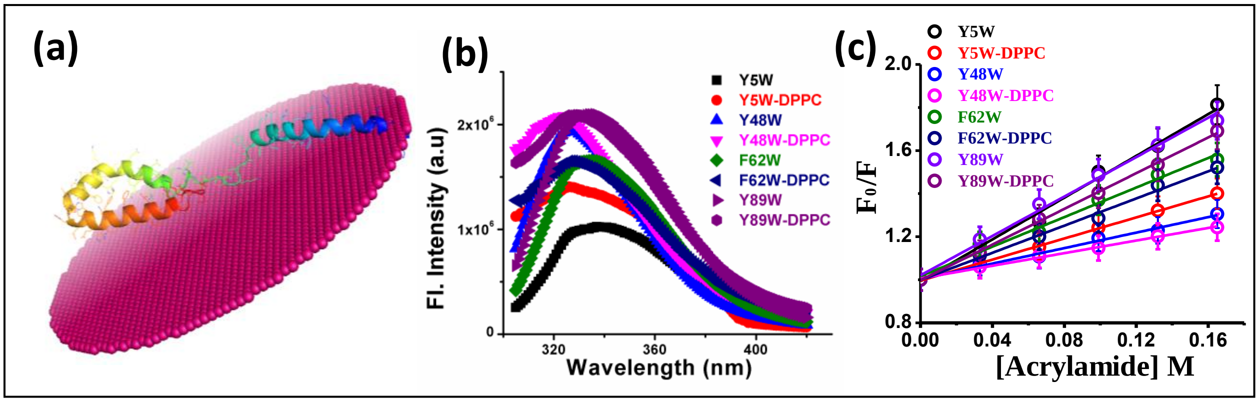


**Figure S11:Detection of the binding region of KMP-11 with membrane.** (a) Positioning of KMP-11 on bilayer structure as monitored by OPM prediction and showed N-targeting nature of KMP-11. (b)Tryptophan fluorescence spectra of four mutants in absence and presence of DPPC model membrane. (c) Acrylamide quenching plots ($F_0/F$ vs acrylamide concentrations) of four mutants i.e Y5W, Y48W, F62W and Y89W in membrane bound and unbound conditions.

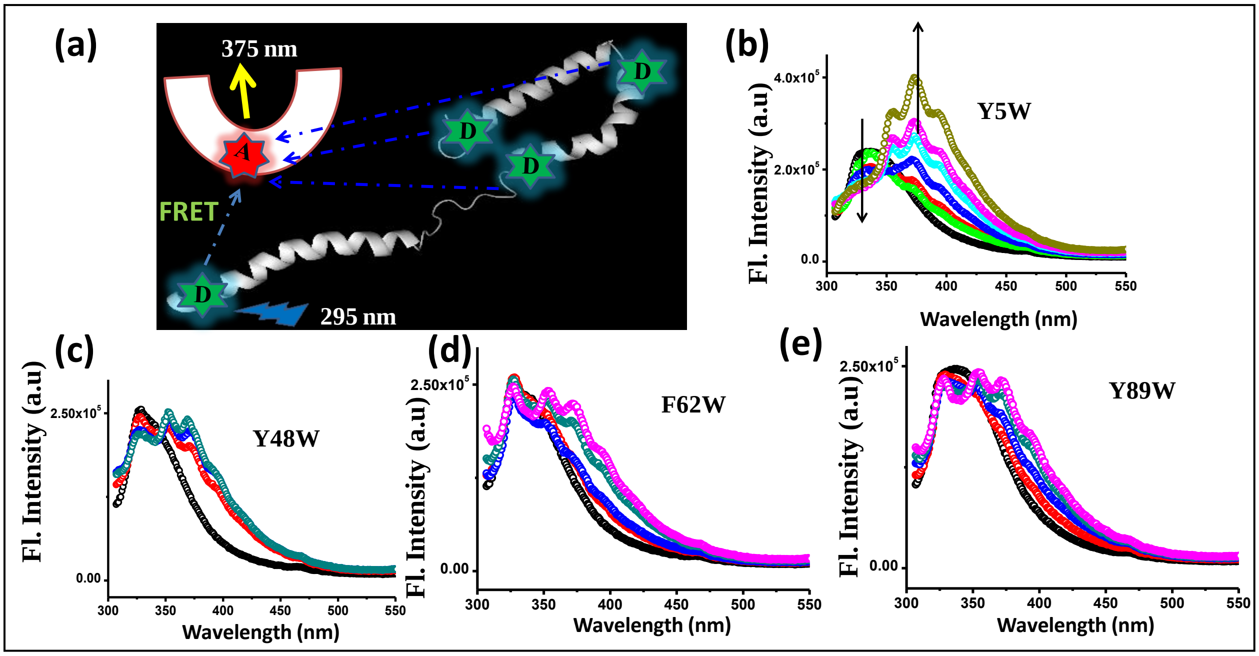


**Figure S12:Energy transfer study of four single tryptophan mutants of KMP-11.** (a) Representative FRET pair was shown where tryptophans at different positions act as donor and DHE inside the bilayer behaved as acceptor.Energy transfer study of the four single tryptophan mutants considering tryptophan residue as donor and membrane bound DHE as acceptor at pH 7.5 [(b) Y5W; (c) Y48W; (d) F62W; (e) Y89W]. The donor acceptor distance and efficiency of energy transfer have been calculated.

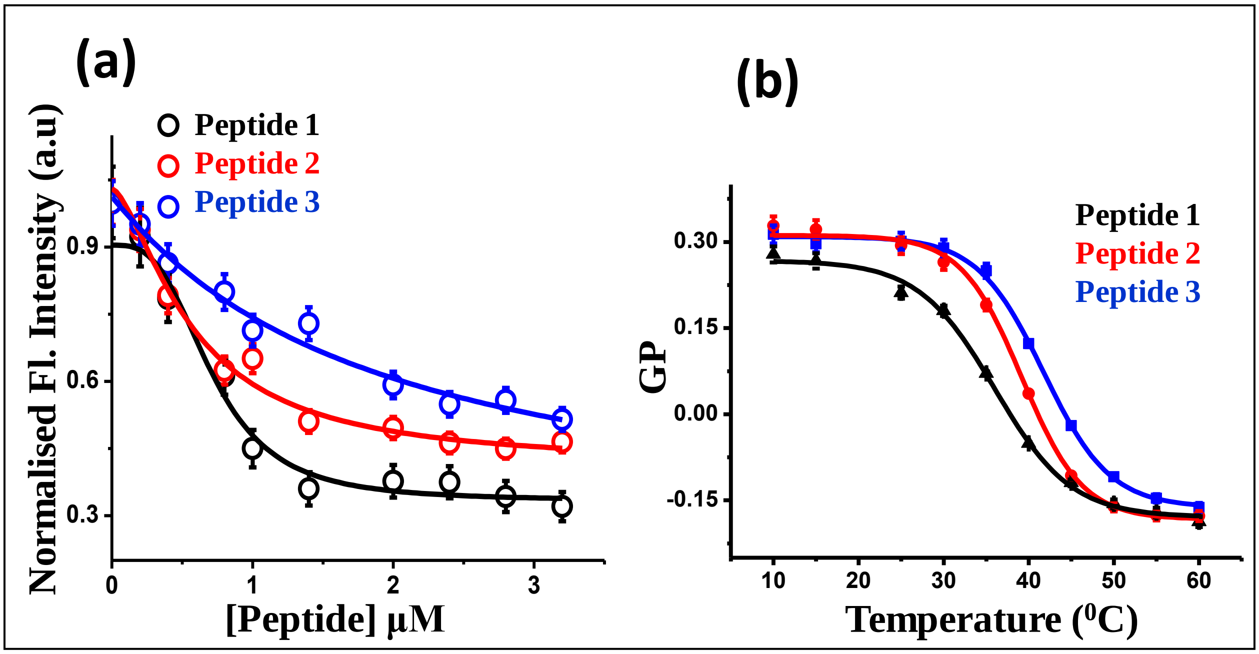


**Figure S13: Binding study of three different peptides with phospholipid model membrane.** (a) Binding of three different peptides with DPPC model membranes as monitored by decrease of DiI C-18 fluorescence intensity with increasing concentration of added peptides. The solid lines come from the fitting of the data using Hill equation. (b) Laurdan generalized polarization of DPPC membrane in presence of different peptides. Typical peptide concentration was chosen 50 µM.

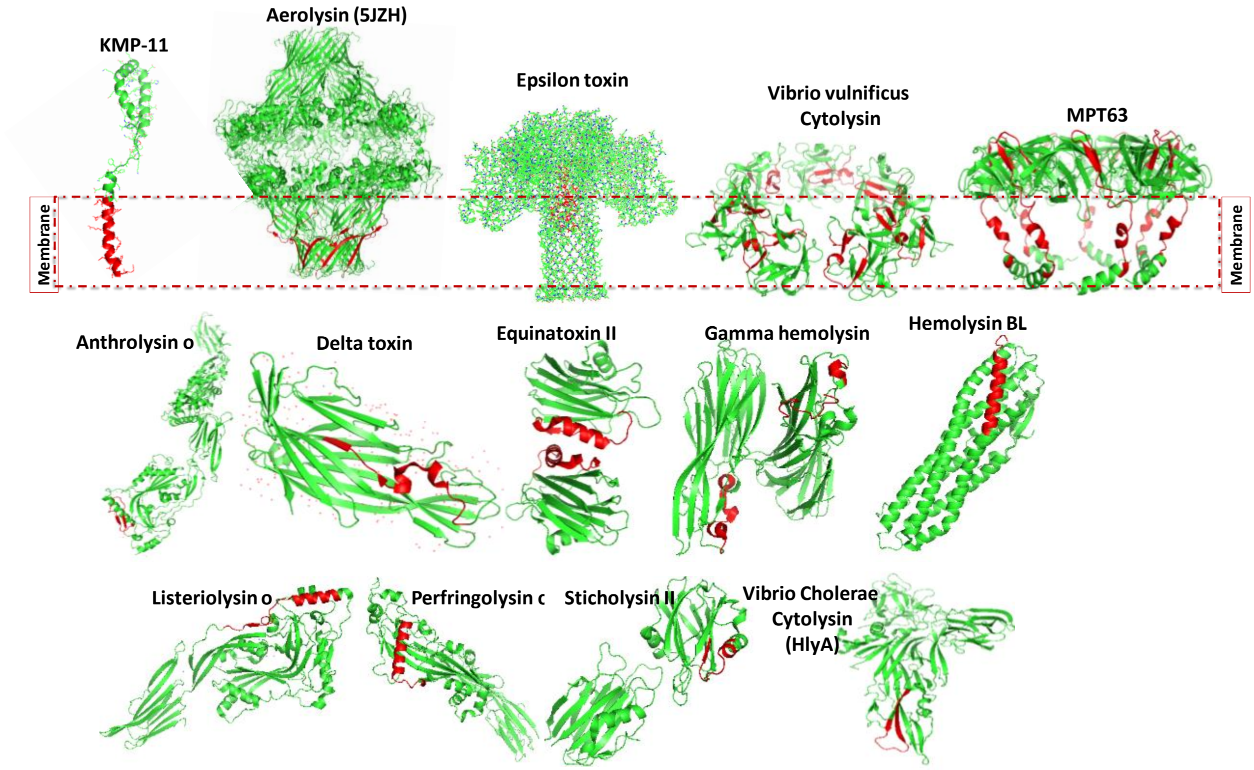


**Figure S14: The sequence similarity between KMP-11-membrane interacting region with the membrane interacting stretches of other disease promoting proteins.** It was found that N-terminal region of KMP-11 remained conserved in all the other proteins(these conserved sequences were marked red). Interestingly, the sequences in other disease promoting proteins which were symmetrical to the KMP-11-N-terminal stretch were found to be positioned in the pore forming regions of the disease proteins.

```
sp|Q36736.1|KM11_LEIDO          ------------------------------------------------------------   0
5JZH:A|PDBID|CHAIN|SEQUENCE     AEPVYPDQLRLFSLGQGVCGDKYRPVNREEAQSVKSNIVGMMGQWQISGLANGWVIMGPG   60

sp|Q36736.1|KM11_LEIDO          ------------------------------------------------------------   0
5JZH:A|PDBID|CHAIN|SEQUENCE     YNGEIKPGTASNTWCYPTNPVTGEIPTLSALDIPDGDEVDVQWRLVHDSANFIKPTSYLA   120

sp|Q36736.1|KM11_LEIDO          -----------------------------------------------MATTYEEFSAKLD   13
5JZH:A|PDBID|CHAIN|SEQUENCE     HYLGYAWVGGNHSQYVGEDMDVTRDGDGWVIRGNNDGGCDGYRCGDKTAIKVSNFAYNLD   180
                                                                                * . .:*: :**

sp|Q36736.1|KM11_LEIDO          RLDQEF------NRKMQEQNAKFFADKPDE------STLSPEMREH------YEKFERMI   55
5JZH:A|PDBID|CHAIN|SEQUENCE     PDSFKHGDVTQSDRQLVKTVVGWAVNDSDTPQSGYDVTLRGDTATNWSKTNTYGLSEKVT   240
                                  . :.      :*:: :  . : .:. *        **  :   :      *   *::

sp|Q36736.1|KM11_LEIDO          KEHTEKFNKK------------------------------------------MHEHSEHF   73
5JZH:A|PDBID|CHAIN|SEQUENCE     TK--NKFKWPLVGETELSIEIAANQSWASQNGGSTTTSLSQSVRPTVPARSKIPVKIELY   298
                                .:  :**:                                            :  : * :

sp|Q36736.1|KM11_LEIDO          KQK----------------FAELLEQQKAAQY----------------------------   89
5JZH:A|PDBID|CHAIN|SEQUENCE     KADISYPYEFKADVSYDLTLSGFLRWGGNAWYTHPDNRPNWNHTFVIGPYKDKASSIRYQ   358
                                * .                ::  :*.    * *

sp|Q36736.1|KM11_LEIDO          ------PSK---------------------------------------------------   92
5JZH:A|PDBID|CHAIN|SEQUENCE     WDKRYIPGEVKWWDWNWTIQQNGLSTMQNNLARVLRPVRAGITGDFSAESQFAGNIEIGA   418
                                      *.:

sp|Q36736.1|KM11_LEIDO          ------ 92
5JZH:A|PDBID|CHAIN|SEQUENCE     PVPLAA 424
```

**Figure S15: Sequence alignment of KMP-11 and Aerolysin (PDB ID: 5JZH) showed 12.73% sequence similarity.**

```
sp|Q36736.1|KM11_LEIDO          -------------------------------------------------MATTYEEFSAK   11
1CA1:A|PDBID|CHAIN|SEQUENCE     WDGKIDGTGTHAMIVTQGVSILENDLSKNEPESVRKNLEILKENMHELQLGSTYPDYDKN   60
                                                                                 :.:** ::. :

sp|Q36736.1|KM11_LEIDO          LDRLDQ-----------------------------------------EFNRKMQE-----   25
1CA1:A|PDBID|CHAIN|SEQUENCE     AYDLYQDHFWDPDTDNNFSKDNSWYLAYSIPDTGESQIRKFSALARYEWQRGNYKQATFY   120
                                   * *                                         *::*   :

sp|Q36736.1|KM11_LEIDO          --QNAKFFADKPDE--S-TLSPEMREHYEKFERMIKEHTEKFNKKMHEHSEHFKQKFAEL   80
1CA1:A|PDBID|CHAIN|SEQUENCE     LGEAMHYFGDIDTPYHPANVTAVDSAGHVKFETFAEERKEQYKINTVG-CKTNEDFYADI   179
                                  :  ::*.*        .::      : *** : :*:.*::: :    .:  :: :*::

sp|Q36736.1|KM11_LEIDO          LEQQKA------------------------------------------------------   86
1CA1:A|PDBID|CHAIN|SEQUENCE     LKNKDFNAWSKEYARGFAKTGKSIYYSHASMSHSWDDWDYAAKVTLANSQKGTAGYIYRF   239
                                *:::.

sp|Q36736.1|KM11_LEIDO          ------------------------------------------------------------   86
1CA1:A|PDBID|CHAIN|SEQUENCE     LHDVSEGNDPSVGKNVKELVAYISTSGEKDAGTDDYMYFGIKTKDGKTQEWEMDNPGNDF   299

sp|Q36736.1|KM11_LEIDO          --------------------------------AQYPSK----------------------   92
1CA1:A|PDBID|CHAIN|SEQUENCE     MTGSKDTYTFKLKDENLKIDDIQNMWIRKRKYTAFPDAYKPENIKVIANGKVVVDKDINE   359
                                                                : :*.

sp|Q36736.1|KM11_LEIDO          -----------    92
1CA1:A|PDBID|CHAIN|SEQUENCE     WISGNSTYNIK    370
```

**Figure S16: Sequence alignment of KMP-11 and Alpha toxin from C. perfringens (PDB ID: 1CA1) showed 14.5% sequence similarity.**

```
sp|Q36736.1|KM11_LEIDO          ------------------------------------------------------------   0
3CQF:A|PDBID|CHAIN|SEQUENCE     MHHHHHHAAAMETQAGNATGAIKNASDINTGIANLKYDSRDILAVNGDKVESFIPKESIN   60
3CQF:B|PDBID|CHAIN|SEQUENCE     MHHHHHHAAAMETQAGNATGAIKNASDINTGIANLKYDSRDILAVNGDKVESFIPKESIN   60


sp|Q36736.1|KM11_LEIDO          ------------------------------------------------------------   0
3CQF:A|PDBID|CHAIN|SEQUENCE     SNGKFVVVEREKKSLTTSPVDILIIDSVVNRTYPGAVQLANKAFADNQPSLLVAKRKPLN   120
3CQF:B|PDBID|CHAIN|SEQUENCE     SNGKFVVVEREKKSLTTSPVDILIIDSVVNRTYPGAVQLANKAFADNQPSLLVAKRKPLN   120


sp|Q36736.1|KM11_LEIDO          ----------------MATTYEEFSAKLDRLDQEFNRKMQEQ------------------   26
3CQF:A|PDBID|CHAIN|SEQUENCE     ISIDLPGMRKENTITVQNPTYGNVAGAVDDLVSTWNEKYSTTHTLPARMQYTESMVYSKS   180
3CQF:B|PDBID|CHAIN|SEQUENCE     ISIDLPGMRKENTITVQNPTYGNVAGAVDDLVSTWNEKYSTTHTLPARMQYTESMVYSKS   180
                                                   ** :.:. :* * . :*.* .

sp|Q36736.1|KM11_LEIDO          --------NAK------------------------------------------FFADKP-   35
3CQF:A|PDBID|CHAIN|SEQUENCE     QIASALNVNAKYLDNSLNIDFNAVANGEKKVMVAAYKQIFYTVSAELPNNPSDLFDNSVT   240
3CQF:B|PDBID|CHAIN|SEQUENCE     QIASALNVNAKYLDNSLNIDFNAVANGEKKVMVAAYKQIFYTVSAELPNNPSDLFDNSVT   240
                                        ***                                          :* :.

sp|Q36736.1|KM11_LEIDO          --------DESTLSPEM-------REHYEKFE-------------RMIKEHTE-------   60
3CQF:A|PDBID|CHAIN|SEQUENCE     FDELTRKGVSNSAPPVMVSNVAYGRTVYVKLETTSKSKDVQAAFKALLKNNSVETSGQYK   300
3CQF:B|PDBID|CHAIN|SEQUENCE     FDELTRKGVSNSAPPVMVSNVAYGRTVYVKLETTSKSKDVQAAFKALLKNNSVETSGQYK   300
                                         ..:  * *       *  * *:*              ::*:::

sp|Q36736.1|KM11_LEIDO          -----------KFNKKMHEHSEHFKQKFAELLEQ--------QKAAQYPSK---------   92
3CQF:A|PDBID|CHAIN|SEQUENCE     DIFEESTFTAVVLGGDAKEHNKVVTKDFNEIRNIIKDNAELSFKNPAYPISYTSTFLKDN   360
3CQF:B|PDBID|CHAIN|SEQUENCE     DIFEESTFTAVVLGGDAKEHNKVVTKDFNEIRNIIKDNAELSFKNPAYPISYTSTFLKDN   360
                                            :. . :**.: ..:.* *: :          *   ** .

sp|Q36736.1|KM11_LEIDO          ------------------------------------------------------------   92
3CQF:A|PDBID|CHAIN|SEQUENCE     ATAAVHNNTDYIETTTTEYSSAKMTLDHYGAYVAQFDVSWDEFTFDQNGKEVLTHKTWEG   420
3CQF:B|PDBID|CHAIN|SEQUENCE     ATAAVHNNTDYIETTTTEYSSAKMTLDHYGAYVAQFDVSWDEFTFDQNGKEVLTHKTWEG   420


sp|Q36736.1|KM11_LEIDO          ------------------------------------------------------------   92
3CQF:A|PDBID|CHAIN|SEQUENCE     SGKDKTAHYSTVIPLPPNSKNIKIVARECTGLAWEWWRTIINEQNVPLTNEIKVSIGGTT   480
3CQF:B|PDBID|CHAIN|SEQUENCE     SGKDKTAHYSTVIPLPPNSKNIKIVARECTGLAWEWWRTIINEQNVPLTNEIKVSIGGTT   480


sp|Q36736.1|KM11_LEIDO          ---------      92
3CQF:A|PDBID|CHAIN|SEQUENCE     LYPTATISH      489
3CQF:B|PDBID|CHAIN|SEQUENCE     LYPTATISH      489
```

**Figure S17: Sequence alignment of KMP-11 and Anthrolysin O (PDB ID: 3CQF) showed 11.45% similarity.**

```
sp|Q36736.1|KM11_LEIDO          -----------------------------MATT---------------------------    4
2XH6:A|PDBID|CHAIN|SEQUENCE     MLSNNLNPMVFENAKEVFLISEDLKTPINITNSNSNLSDGLYVIDKGDGWILGEPSVVSS    60
2XH6:B|PDBID|CHAIN|SEQUENCE     MLSNNLNPMVFENAKEVFLISEDLKTPINITNSNSNLSDGLYVIDKGDGWILGEPSVVSS    60
2XH6:C|PDBID|CHAIN|SEQUENCE     MLSNNLNPMVFENAKEVFLISEDLKTPINITNSNSNLSDGLYVIDKGDGWILGEPSVVSS    60
                                                             ::.:

sp|Q36736.1|KM11_LEIDO          -------YEEFSAKLD--------------------R--LDQEFNRKMQEQNAKFFADKP    35
2XH6:A|PDBID|CHAIN|SEQUENCE     QILNPNETGTFSQSLTKSKEVSINVNFSVGFTSEFIQASVEYGFGITIGEQNTT---ERS    117
2XH6:B|PDBID|CHAIN|SEQUENCE     QILNPNETGTFSQSLTKSKEVSINVNFSVGFTSEFIQASVEYGFGITIGEQNTT---ERS    117
2XH6:C|PDBID|CHAIN|SEQUENCE     QILNPNETGTFSQSLTKSKEVSINVNFSVGFTSEFIQASVEYGFGITIGEQNTT---ERS    117
                                          ** .*                     :  ::  *. .: ***:.   ::

sp|Q36736.1|KM11_LEIDO          DESTLSPEMR------EHYEKFERMIKEH-------------------------------    58
2XH6:A|PDBID|CHAIN|SEQUENCE     VSTTAGPNEYVYYKVYATYRKYQAIRISHGNISDDGSIYKLTGIWLSKTSADSLGNIDQG    177
2XH6:B|PDBID|CHAIN|SEQUENCE     VSTTAGPNEYVYYKVYATYRKYQAIRISHGNISDDGSIYKLTGIWLSKTSADSLGNIDQG    177
2XH6:C|PDBID|CHAIN|SEQUENCE     VSTTAGPNEYVYYKVYATYRKYQAIRISHGNISDDGSIYKLTGIWLSKTSADSLGNIDQG    177
                                 .:* .*:          *.*:: :  .*

sp|Q36736.1|KM11_LEIDO          ----------------------------TEKFN-------------------------KK    65
2XH6:A|PDBID|CHAIN|SEQUENCE     SLIETGERCVLTVPSTDIEKEILDLAAATERLNLTDALNSNPAGNLYDWRSSNSYPWTQK    237
2XH6:B|PDBID|CHAIN|SEQUENCE     SLIETGERCVLTVPSTDIEKEILDLAAATERLNLTDALNSNPAGNLYDWRSSNSYPWTQK    237
2XH6:C|PDBID|CHAIN|SEQUENCE     SLIETGERCVLTVPSTDIEKEILDLAAATERLNLTDALNSNPAGNLYDWRSSNSYPWTQK    237
                                                            **::*                         :*

sp|Q36736.1|KM11_LEIDO          MHEH-------------------SEHFKQKFAELLEQQ----------------------    84
2XH6:A|PDBID|CHAIN|SEQUENCE     LNLHLTITATGQKYRILASKIVDFNIYSNNFNNLVKLEQSLGDGVKDHYVDISLDAGQYV    297
2XH6:B|PDBID|CHAIN|SEQUENCE     LNLHLTITATGQKYRILASKIVDFNIYSNNFNNLVKLEQSLGDGVKDHYVDISLDAGQYV    297
2XH6:C|PDBID|CHAIN|SEQUENCE     LNLHLTITATGQKYRILASKIVDFNIYSNNFNNLVKLEQSLGDGVKDHYVDISLDAGQYV    297
                                :: *                    : :.::* :*:: :

sp|Q36736.1|KM11_LEIDO          --------KAAQYPSK------ 92
2XH6:A|PDBID|CHAIN|SEQUENCE     LVMKANSSYSGNYPYSILFQKF 319
2XH6:B|PDBID|CHAIN|SEQUENCE     LVMKANSSYSGNYPYSILFQKF 319
2XH6:C|PDBID|CHAIN|SEQUENCE     LVMKANSSYSGNYPYSILFQKF 319
                                         :.:** .
```

**Figure S18: Sequence alignment of KMP-11 and Clostridium perfringens enterotoxin (PDB ID: 2XH6) showed 18.80% similarity.**

```
CLUSTAL O(1.2.4) multiple sequence alignment

1COL:A|PDBID|CHAIN|SEQUENCE      VAE--------KAK----DERELLEKTSELIAGMGD--KIGEHLGDKYKAIAKDIADNIK   46
sp|Q36736.1|KM11_LEIDO           MATTYEEFSAKLDRLDQEFNRKMQEQNAKFFADKPDESTLSPEMREHYEKFERMIKEHTE   60
                                 :*           :     :*:: *:.::::*.  *  .:. .: ::*: : : * :: :

1COL:A|PDBID|CHAIN|SEQUENCE      NFQGKTIRSFDDAMASLNKITANPAMKINKADRDALVNAWKHVDAQDMANKLGNLSKAFK   106
sp|Q36736.1|KM11_LEIDO           KFNKKM-----------------------------------HEHSEHFKQKFAELLEQQK   85
                                 :*: *                                    * .::.: :*:.:* :  *

1COL:A|PDBID|CHAIN|SEQUENCE      VADVVMKVEKVREKSIEGYETGNWGPLMLEVESWVLSGIASSVALGIFSATLGAYALSLG   166
sp|Q36736.1|KM11_LEIDO           A---------------AQYPSK--------------------------------------   92
                                 .                 * :

1COL:A|PDBID|CHAIN|SEQUENCE      VPAIAVGIAGILLAAVVGALIDDKFADALNNEIIRPAH 204
sp|Q36736.1|KM11_LEIDO           -------------------------------------- 92
```

**Figure S19: Sequence alignment of KMP-11 and colicin A (PDB ID: 1COL) showed 27.45% similarity.**

```
sp|Q36736.1|KM11_LEIDO          ------------------------------------------------------------   0
2YGT:A|PDBID|CHAIN|SEQUENCE     GSNDLGSKSEIRKEENGNVTIITQNNKQIRKYSSTDSATTKSNSKITVDASFVDDKFSSE   60

sp|Q36736.1|KM11_LEIDO          ------------------------------------------------------------   0
2YGT:A|PDBID|CHAIN|SEQUENCE     MTTIISLKGFIPSGRKIFALSKYRGVMRWPIKYMVDLKNNSLDSSVKIVDSVPKNTISTK   120

sp|Q36736.1|KM11_LEIDO          ------------------------------------------------------------   0
2YGT:A|PDBID|CHAIN|SEQUENCE     EVNNTISYSIGGGIDTSNKASLNANYAVSKSISYVQPDYNTIQTNDTNSIASWNTEFAET   180

sp|Q36736.1|KM11_LEIDO          ------------------MATTYEE-------------------FSAKL-----------   12
2YGT:A|PDBID|CHAIN|SEQUENCE     RDGYNVNSWNIVYGNQMFMRSRYSGTSTTNFTPDYQLSSLITGGFSPNFGVVLTAPNGTK   240
                                                  * : *.                    ** ::

sp|Q36736.1|KM11_LEIDO          -DRLDQEFNRKMQEQN--------AKFFADKPDESTLSPEMREHYEKFERMIKEHTEKFN   63
2YGT:A|PDBID|CHAIN|SEQUENCE     KSQIEISLKREINSYHIAWDTEWQGRNYPDSKIEET---------VKFELDWEKHTIR--   289
                                 .::: .::*:::. :        .: : *.  *.*          ***   ::** :

sp|Q36736.1|KM11_LEIDO          KKMHEHSEHFKQKFAELLEQQKAAQYPSK  92
2YGT:A|PDBID|CHAIN|SEQUENCE     QISHHHHHH--------------------  298
                                :  *.* .*
```

**Figure S20: Sequence alignment of KMP-11 and Delta toxin (PDB ID: 2YGT) showed 15.43% similarity.**

```
CLUSTAL O(1.2.4) multiple sequence alignment


sp|Q36736.1|KM11_LEIDO             ------------------------MATTYEEFSA------------KLDRLDQEFNR---    21
6RB9:A|PDBID|CHAIN|SEQUENCE        MSYYHHHHHHDYDIPTTENLYFQGAMASYDNVDTLIEKGRYNTKYNYLKRMEKYYPNAMA    60
                                                             ::*::..:             *.*::: : .

sp|Q36736.1|KM11_LEIDO             -----KMQEQNAKFFADKPDEST-LSPEMREHYEKFE--RMIKEH---------------    58
6RB9:A|PDBID|CHAIN|SEQUENCE        YFDKVTINPQGNDFYINNPKVELDGEPSMNYLEDVYVGKALLTNDTQQEQKLKSQSFTCK    120
                                        .:: *. .*: ::*. .   .*.*.   : :    ::.:.

sp|Q36736.1|KM11_LEIDO             -------------------TEKFNKKMHEHSEHFKQK--FA-------------------    78
6RB9:A|PDBID|CHAIN|SEQUENCE        NTDTVTATTTHTVGTSIQATAKFTVPFNETGVSLTTSYSFANTNTNTNSKEITHNVPSQD    180
                                                      * **.  ::* .  :. .  **

sp|Q36736.1|KM11_LEIDO             ------------------------ELLEQQKAA---------QYPSK-------------    92
6RB9:A|PDBID|CHAIN|SEQUENCE        ILVPANTTVEVIAYLKKVNVKGNVKLVGQVSGSEWGEIPSYLAFPRDGYKFSLSDTVNKS    240
                                                           :*: * ..:          :* .

sp|Q36736.1|KM11_LEIDO             ---------------------------------------------- 92
6RB9:A|PDBID|CHAIN|SEQUENCE        DLNEDGTININGKGNYSAVMGDELIVKVRNLNTNNVQEYVIPVDKK 286
```

**Figure S21: Sequence alignment of KMP-11 and Epsilon toxin (PDB ID: 6RB9) showed 22.02% similarity.**

```
sp|Q36736.1|KM11_LEIDO          -----------MATTYEEFSAKLDRLD---QE----------------------------    18
1IAZ:A|PDBID|CHAIN|SEQUENCE     SADVAGAVIDGASLSFDILKTVLEALGNVKRKIAVGVDNESGKTWTALNTYFRSGTSDIV    60
1IAZ:B|PDBID|CHAIN|SEQUENCE     SADVAGAVIDGASLSFDILKTVLEALGNVKRKIAVGVDNESGKTWTALNTYFRSGTSDIV    60
                                            : ::: :.: *: *.   ::

sp|Q36736.1|KM11_LEIDO          FNRKMQEQNAKFFADKPDESTLSP------------------EM--REHYEKFERMIKEH    58
1IAZ:A|PDBID|CHAIN|SEQUENCE     LPHKVPHGKALLYNGQKDRGPVATGAVGVLAYLMSDGNTLAVLFSVPYDYNWYSNWWNV-    119
1IAZ:B|PDBID|CHAIN|SEQUENCE     LPHKVPHGKALLYNGQKDRGPVATGAVGVLAYLMSDGNTLAVLFSVPYDYNWYSNWWNV-    119
                                : :*: . :* :: .: *.. ::                    :    .*: :..  :

sp|Q36736.1|KM11_LEIDO          TEKFNKKMHEHSEHFKQKFAELLEQQKAAQYPSK--------------------------    92
1IAZ:A|PDBID|CHAIN|SEQUENCE     -RIYKGKRRADQRMYEELYYNLSPFRGDNGWHTRNLGYGLKSRGFMNSSGHAILEIHVSK    178
1IAZ:B|PDBID|CHAIN|SEQUENCE     -RIYKGKRRADQRMYEELYYNLSPFRGDNGWHTRNLGYGLKSRGFMNSSGHAILEIHVSK    178
                                 . :: * : ... ::: : :*   :    : ::

sp|Q36736.1|KM11_LEIDO          -      92
1IAZ:A|PDBID|CHAIN|SEQUENCE     A      179
1IAZ:B|PDBID|CHAIN|SEQUENCE     A      179
```

**Figure S22: Sequence alignment of KMP-11 and Equinatoxin II (PDB ID:1IAZ) showed 30% similarity.**

```
sp|Q36736.1|KM11_LEIDO            ------------------------------------------------------------    0
2QK7:A|PDBID|CHAIN|SEQUENCE       GPLGSPEFENKIEDIGQ-----GAEIIKRTQDITSKRLAICQNIQFDFVKDKKYNKDALV    55
2QK7:B|PDBID|CHAIN|SEQUENCE       GPLGSPEFEGKITPVSVKKVDDKVTLYKTTATADSDKFKISQILTFNFIKDKSYDKDTLV    60

sp|Q36736.1|KM11_LEIDO            ------------------------------------------------------------    0
2QK7:A|PDBID|CHAIN|SEQUENCE       VKMQGFISSRTTYSDLKKYPYIKRMIWPFQYNISLKTK-DSNVDLINYLPKNKIDSADVS    114
2QK7:B|PDBID|CHAIN|SEQUENCE       LKAAGNINSGYEKPNPNDYDF-SKLYWGAKYNVSISSQSNDSVNVVDYAPKNQNEEFQVQ    119

sp|Q36736.1|KM11_LEIDO            ------------------------------------------------------------    0
2QK7:A|PDBID|CHAIN|SEQUENCE       QKLGYNIGGNFQSAPSIGG--SGSFNYSKTISYNQKNYVTEVE-SQNSKGVKWGVKANSF    171
2QK7:B|PDBID|CHAIN|SEQUENCE       NTLGYTFGGDISISNGLSGGLNGNTAFSETINYKQESYRTTLSRCTNYKNVGWGVEAHKI    179

sp|Q36736.1|KM11_LEIDO            ------------MATTYEEFS---------------------------------------    9
2QK7:A|PDBID|CHAIN|SEQUENCE       VTPN----GQV---SAYDQYL-FAQDPTGPAARDYFVPDNQLPPLIQSGFNPSFITTLSH    223
2QK7:B|PDBID|CHAIN|SEQUENCE       MNNGWGPYGRDSFHPTYGNELFLAGRQSSAYAGQNFIAQHQMPLLSRSNFNPEFLSVLSH    239
                                                 :* :

sp|Q36736.1|KM11_LEIDO            ----AKLDRLDQEFNRKMQEQNAK----F-FADKPDES----TLSPEMREHYEKFERMIK    56
2QK7:A|PDBID|CHAIN|SEQUENCE       EKGKGDKSEFEITYGRNMDATYAYVTRHRLAVDRKHDAFKNRNVTVKYEVNWKTHEVKIK    283
2QK7:B|PDBID|CHAIN|SEQUENCE       RQDGAKKSKITVTYQREMDLYQIRWNG-FYWAGANYKNFKTRTFKSTYEIDWENHKVKLL    298
                                      .. ..:   : *:*:            ..   .     ...   . .::..:  :

sp|Q36736.1|KM11_LEIDO            EHTEKFNKKMHEHSEHFKQKFAELLEQQKAAQYPSK    92
2QK7:A|PDBID|CHAIN|SEQUENCE       SITPK-------------------------------    288
2QK7:B|PDBID|CHAIN|SEQUENCE       DTKETENNK---------------------------    307
                                  . . .
```

**Figure S23: Sequence alignment of KMP-11 and Gama hemolysin (PDB ID:2QK7) showed 10.09 % similarity.**

```
sp|Q36736.1|KM11_LEIDO          ------------------------------------------------------MATTYE    6
2NRJ:A|PDBID|CHAIN|SEQUENCE     SLSEIEQTNNGDTALSANEARMKETLQKAGLFAKSMNAYSYMLIKNPDVNFEGITINGYV    60
                                                                                        . *

sp|Q36736.1|KM11_LEIDO          EFSAKLDRLD-------QEFNRKMQEQN--------------AKFFA------DKPDEST    39
2NRJ:A|PDBID|CHAIN|SEQUENCE     DLPGRIVQDQKNARAHAVTWDTKVKKQLLDTLNGIVEYDTTFDNYYETMVEAINTGDGET    120
                                :: .:: : :         :: *:::*                :::       :. * .*

sp|Q36736.1|KM11_LEIDO          LS-------PEMREHYEKFERMIKEHTEKFNKKMHEHSEHFKQ---KFAELLEQQKA---    86
2NRJ:A|PDBID|CHAIN|SEQUENCE     LKEGITDLRGEIQQN-QKYAQQLIEELTKLRDSIGHDVRAFGSNKELLQSILKNQGADVD    179
                                *.        *:::: :*: : : *.  *:...: .. . * .    : .:*::* *

sp|Q36736.1|KM11_LEIDO          ------------AQY---------------------------------------------    89
2NRJ:A|PDBID|CHAIN|SEQUENCE     ADQKRLEEVLGSVNYYKQLESDGFNVMKGAILGLPIIGGIIVGVARDNLGKLEPLLAELR    239
                                            .:*

sp|Q36736.1|KM11_LEIDO          ------------------------------------------------------------    89
2NRJ:A|PDBID|CHAIN|SEQUENCE     QTVDYKVTLNRVVGVAYSNINEMHKALDDAINALTYMSTQWHDLDSQYSGVLGHIENAAQ    299

sp|Q36736.1|KM11_LEIDO          --------------------------------------------PSK        92
2NRJ:A|PDBID|CHAIN|SEQUENCE     KADQNKFKFLKPNLNAAKDSWKTLRTDAVTLKEGIKELKVETVTPQK        346
                                                                            *.*
```

**Figure S24: Sequence alignment of KMP-11 and Hemolysin BL (PDB ID:2NRJ) showed 17.91% similarity.**

```
CLUSTAL O(1.2.4) multiple sequence alignment


sp|Q36736.1|KM11_LEIDO          MATTYEE----------------------------------------FSA--KLDRLDQE   18
1W3G:A|PDBID|CHAIN|SEQUENCE     MTDIYIPPEGLYFRLLGFASRQVIFARNSPSPDVGLSPVNDQATDQYFSLIYGTGEHAGL   60
                                *:  *                                          **     ..

sp|Q36736.1|KM11_LEIDO          FNRKMQEQNAKFFADKPDESTLSPEM---------------REHYEKFERMIKEHTEK-F   62
1W3G:A|PDBID|CHAIN|SEQUENCE     YAIKSKATGKVLFSRRPAEPYVGQIDGDGRYPDNWFKIEPGKTYLSKYFRLVQPSTGTAL   120
                                :  * :  .  :*: :* *  :.                  : : .*: *:::  * . :

sp|Q36736.1|KM11_LEIDO          NKKM--------HEHSEHFKQKFAE-----------------------------------   79
1W3G:A|PDBID|CHAIN|SEQUENCE     VSRTHLQPYFWNHPQTEVFDDQYFTFLFEDMSIDKIEYDLKDGRILSSTPNVLATQTLEN   180
                                 .:         * ::* *.:::

sp|Q36736.1|KM11_LEIDO          --------------LLEQQKAAQYPSK---------------------------------   92
1W3G:A|PDBID|CHAIN|SEQUENCE     TSSQTQEMSFNLSQTLTQTSTFAYTAGFTIAVGTAFKAGVPIFAETEFKVDISVDNQWNW   240
                                               * * .:  * :

sp|Q36736.1|KM11_LEIDO          ------------------------------------------------------------   92
1W3G:A|PDBID|CHAIN|SEQUENCE     GEENTFSKTYTATFSVRAGPGETVKAVSTVDSGIINVPFTAYLSSKSTGFEVTTEGIWRG   300


sp|Q36736.1|KM11_LEIDO          ---------------       92
1W3G:A|PDBID|CHAIN|SEQUENCE     VSSWDLRHTLTSVTA       315
```

**Figure S25: Sequence alignment of KMP-11 and Hemolysin lectin (PDB ID:1W3G) showed 15.23 % similarity.**

```
sp|Q36736.1|KM11_LEIDO          -----------MAT----------------------------------------------    3
1S3R:A|PDBID|CHAIN|SEQUENCE     MGGSHHHHHHGMASMTGGQQMGRDLYDDDDKDRWGSETPTKPKAAQTEKKTEKKPENSNS   60
1S3R:B|PDBID|CHAIN|SEQUENCE     MGGSHHHHHHGMASMTGGQQMGRDLYDDDDKDRWGSETPTKPKAAQTEKKTEKKPENSNS   60
                                           **:

sp|Q36736.1|KM11_LEIDO          --------------TYEE------------------------------------------    7
1S3R:A|PDBID|CHAIN|SEQUENCE     EAAKKALNDYIWGLQYDKLNILTHQGEKLKNHSSREAFHRPGEYVVIEKKKQSISNATSK   120
1S3R:B|PDBID|CHAIN|SEQUENCE     EAAKKALNDYIWGLQYDKLNILTHQGEKLKNHSSREAFHRPGEYVVIEKKKQSISNATSK   120
                                               *::

sp|Q36736.1|KM11_LEIDO          -----------FSAKLDRLDQEFNRK--------------------MQEQNAKFFADKPD   36
1S3R:A|PDBID|CHAIN|SEQUENCE     LSVSSANDDRIFPGALLKADQSLLENLPTLIPVNRGKTTISVNLPGLKNGESNLTVENPS   180
1S3R:B|PDBID|CHAIN|SEQUENCE     LSVSSANDDRIFPGALLKADQSLLENLPTLIPVNRGKTTISVNLPGLKNGESNLTVENPS   180
                                           * . * : **.: .:                    ::: :::: .::*.

sp|Q36736.1|KM11_LEIDO          ESTLSPEMREHYEKFERMI--------------------KEHTEKFNKKMHEH-------   69
1S3R:A|PDBID|CHAIN|SEQUENCE     NSTVRTAVNNLVEKWIQNYSKTHAVPARMQYESISAQSMSQLQAKFGADFSKVGAPLNVD   240
1S3R:B|PDBID|CHAIN|SEQUENCE     NSTVRTAVNNLVEKWIQNYSKTHAVPARMQYESISAQSMSQLQAKFGADFSKVGAPLNVD   240
                                :**:   :.:  **: :                      .:   **. .: :

sp|Q36736.1|KM11_LEIDO          --------SEHFKQKFAELLEQQ--KAAQYPSK---------------------------   92
1S3R:A|PDBID|CHAIN|SEQUENCE     FSSVHKGEKQVFIANFRQVYYTASVDSPNSPSALFGSGITPTDLINRGVNSKTPPVYVSN   300
1S3R:B|PDBID|CHAIN|SEQUENCE     FSSVHKGEKQVFIANFRQVYYTASVDSPNSPSALFGSGITPTDLINRGVNSKTPPVYVSN   300
                                        .: *  :* ::      .: : **

sp|Q36736.1|KM11_LEIDO          ------------------------------------------------------------   92
1S3R:A|PDBID|CHAIN|SEQUENCE     VSYGRAMYVKFETTSKSTKVQAAIDAVVKGAKLKAGTEYENILKNTKITAVVLGGNPGEA   360
1S3R:B|PDBID|CHAIN|SEQUENCE     VSYGRAMYVKFETTSKSTKVQAAIDAVVKGAKLKAGTEYENILKNTKITAVVLGGNPGEA   360

sp|Q36736.1|KM11_LEIDO          ------------------------------------------------------------   92
1S3R:A|PDBID|CHAIN|SEQUENCE     SKVITGNIDTLKDLIQKGSNFSAQSPAVPISYTTSFVKDNSIATIQNNTDYIETKVTSYK   420
1S3R:B|PDBID|CHAIN|SEQUENCE     SKVITGNIDTLKDLIQKGSNFSAQSPAVPISYTTSFVKDNSIATIQNNTDYIETKVTSYK   420

sp|Q36736.1|KM11_LEIDO          ------------------------------------------------------------   92
1S3R:A|PDBID|CHAIN|SEQUENCE     DGALTLNHDGAFVARFYVYWEELGHDADGYETIRSRSWSGNGYNRGAHYSTTLRFKGNVR   480
1S3R:B|PDBID|CHAIN|SEQUENCE     DGALTLNHDGAFVARFYVYWEELGHDADGYETIRSRSWSGNGYNRGAHYSTTLRFKGNVR   480

sp|Q36736.1|KM11_LEIDO          -------------------------------------------------------        92
1S3R:A|PDBID|CHAIN|SEQUENCE     NIRVKVLGATGLAWEPWRLIYSKNDLPLVPQRNISTWGTTLHPQFEDKVVKDNTD        535
1S3R:B|PDBID|CHAIN|SEQUENCE     NIRVKVLGATGLAWEPWRLIYSKNDLPLVPQRNISTWGTTLHPQFEDKVVKDNTD        535
```

**Figure S26: Sequence alignment of KMP-11 and Intermedilysin (PDB ID:1S3R) showed 10.6 % similarity.**

```
sp|Q36736.1|KM11_LEIDO          ------------------------------------MATTYEEFSAKLDRLDQ------E   18
3LE0:A|PDBID|CHAIN|SEQUENCE     EQGNRPVETENIARGKQASQSSTAYGGAATRAVDGNVDSDYGHHSVTHTNFEDNAWWQVD   60
                                                                    : : * ..*..  .:::      :

sp|Q36736.1|KM11_LEIDO          FNRKMQEQNAKFFADKPDESTLSPEMREHYEKFERMIKEHTEKFNKKMHEHSEHFKQKFA   78
3LE0:A|PDBID|CHAIN|SEQUENCE     LGKTENVGKVKLYNRGD--G----NVANRLSNFDVVLLNEAK---------QEVARQHFD   105
                                :.:. :  :.*::      .    :: :: .:*: :: :.::         .*  :*:*

sp|Q36736.1|KM11_LEIDO          EL--------LEQQKAAQYPSK--------------------------       92
3LE0:A|PDBID|CHAIN|SEQUENCE     SLNGKAELEVFFTAKDARYVKVELKTKNTPLSLAEVEVFRSATTQVGC       153
                                .*        :   * *:* .
```

**Figure S27: Sequence alignment of KMP-11 and Lectinolysin (PDB ID:3LE0) showed 32.6% similarity.**

```
sp|Q36736.1|KM11_LEIDO            --------------MATTYEEFS-AKLDRL------D------QEF------NRKMQEQN   27
1LKF:A|PDBID|CHAIN|SEQUENCE       EGKITPVSVKKVDDKVTLYKTTATADSDKFKISQILTFNFIKDKSYDKDTLVLKATGNIN   60
                                                 .* *:  : *. *::             :.:       :   : *

sp|Q36736.1|KM11_LEIDO            AKFFADKPDESTLSPEM-------------REH---------------------------   47
1LKF:A|PDBID|CHAIN|SEQUENCE       SGFVKPNPNDYDFSKLYWGAKYNVSISSQSNDSVNVVDYAPKNQNEEFQVQNTLGYTFGG   120
                                  : *.  :*::  :*                .:

sp|Q36736.1|KM11_LEIDO            --------------YEKFERMIKEHTEKFNKKMH--------------------------   67
1LKF:A|PDBID|CHAIN|SEQUENCE       DISISNGLSGGLNGNTAFSETINYKQESYRTTLSRNTNYKNVGWGVEAHKIMNNGWGPYG   180
                                                   *.. *: : *.:...:

sp|Q36736.1|KM11_LEIDO            -----------------------------------EHSEHFKQKFAELLEQQKAAQYPSK   92
1LKF:A|PDBID|CHAIN|SEQUENCE       RDSFHPTYGNELFLAGRQSSAYAGQNFIAQHQMPLLSRSNFNPEFLSVLSHRQDGAKKSK   240
                                                                        .:*: :* .:*.::: .   **

sp|Q36736.1|KM11_LEIDO            -----------------------------------------------------------    92
1LKF:A|PDBID|CHAIN|SEQUENCE       ITVTYQREMDLYQIRWNGFYWAGANYKNFKTRTFKSTYEIDWENHKVKLLDTKETENNK    299
```

**Figure S28: Sequence alignment of KMP-11 and Leucocidin F (PDB ID:1LKF) showed 18% similarity.**

```
sp|Q36736.1|KM11_LEIDO          ------------------------------------------------------------   0
4CDB:A|PDBID|CHAIN|SEQUENCE     MAPPASPPASPKTPIEKKHADEIDKYIQGLDYNKNNVLVYHGDAVTNVPPRKGYKDGNEY   60

sp|Q36736.1|KM11_LEIDO          ------------------------------------------------------------   0
4CDB:A|PDBID|CHAIN|SEQUENCE     IVVEKKKKSINQNNADIQVVNAISSLTYPGALVKANSELVENQPDVLPVKRDSLTLSIDL   120

sp|Q36736.1|KM11_LEIDO          ----------------------------------MATTYEEFSAKLDRLDQEFNRKMQEQ   26
4CDB:A|PDBID|CHAIN|SEQUENCE     PGMTNQDNKIVVKNATKSNVNNAVNTLVERWNEKYAQAYPNVSAKIDYDDE-----MAYS   175
                                                                   * :* :.***:*  *:     *  .

sp|Q36736.1|KM11_LEIDO          NAKFFADKPDESTLSPEMREHYEKFERMIKEHTEKF--------NKKMHEHSEHFKQKFA   78
4CDB:A|PDBID|CHAIN|SEQUENCE     E----------SQL-------IAKFGTAFKAVNNSLNVNFGAISEGKMQEEVISFKQIYY   218
                                :          * *         **   :*  .:.:        : **:*.   *** :

sp|Q36736.1|KM11_LEIDO          ELLE--QQKAA-----------------------QYPSK---------------------   92
4CDB:A|PDBID|CHAIN|SEQUENCE     NVNVNEPTRPSRFFGKAVTKEQLQALGVNAENPPAYISSVAYGRQVYLKLSTNSHSTKVK   278
                                ::      : :                        * *.

sp|Q36736.1|KM11_LEIDO          ------------------------------------------------------------   92
4CDB:A|PDBID|CHAIN|SEQUENCE     AAFDAAVSGKSVSGDVELTNIIKNSSFKAVIYGGSAKDEVQIIDGNLGDLRDILKKGATF   338

sp|Q36736.1|KM11_LEIDO          ------------------------------------------------------------   92
4CDB:A|PDBID|CHAIN|SEQUENCE     NRETPGVPIAYTTNFLKDNELAVIKNNSEYIETTSKAYTDGKINIDHSGGYVAQFNISWD   398

sp|Q36736.1|KM11_LEIDO          ------------------------------------------------------------   92
4CDB:A|PDBID|CHAIN|SEQUENCE     EVNYDPEGNEIVQHKNWSENNKSKLAHFTSSIYLPGNARNINVYAKECTGLAWEWWRTVI   458

sp|Q36736.1|KM11_LEIDO          ------------------------------ 92
4CDB:A|PDBID|CHAIN|SEQUENCE     DDRNLPLVKNRNISIWGTTLYPKYSNKVDN 488
```

**Figure S29: Sequence alignment of KMP-11 and Listeriolysin O (PDB ID:4CDB) showed 8 % similarity.**

```
sp|Q36736.1|KM11_LEIDO        ------------------------------------------------------------   0
1PFO:A|PDBID|CHAIN|SEQUENCE   MIRFKKTKLIASIAMALCLFSQPVISFSKDITDKNQSIDSGISSLSYNRNEVLASNGDKI   60

sp|Q36736.1|KM11_LEIDO        ------------------------------------------------------------   0
1PFO:A|PDBID|CHAIN|SEQUENCE   ESFVPKEGKKAGNKFIVVERQKRSLTTSPVDISIIDSVNDRTYPGALQLADKALVENRPT   120

sp|Q36736.1|KM11_LEIDO        --------------------------MATTYEEFSAKLDRLDQEFNRKMQEQN-------   27
1PFO:A|PDBID|CHAIN|SEQUENCE   ILMVKRKPININIDLPGLKGENSIKVDDPTYGKVSGAIDELVSKWNEKYSSTHTLPARTQ   180
                                                           ** :.*. :*.* .::*.* .. :

sp|Q36736.1|KM11_LEIDO        ------------------------------------------------------------   27
1PFO:A|PDBID|CHAIN|SEQUENCE   YSESMVYSKSQISSALNVNAKVLENSLGVDFNAVANNEKKVMILAYKQIFYTVSADLPKN   240

sp|Q36736.1|KM11_LEIDO        -AKFFADKPD--------ESTLSPE-M-------REHYE-------------KFERMIKE   57
1PFO:A|PDBID|CHAIN|SEQUENCE   PSDLFDDSVTFNDLKQKGVSNEAPPLMVSNVAYGRTIYVKLETTSSSKDVQAAFKALIKN   300
                               :.:* *.           *. :*  *       *  *               *: :**:

sp|Q36736.1|KM11_LEIDO        HTEK------------------FNKKMHEHSEHFKQKFAELLEQ--------QKAAQYPS   91
1PFO:A|PDBID|CHAIN|SEQUENCE   TDIKNSQQYKDIYENSSFTAVVLGGDAQEHNKVVTKDFDEIRKVIKDNATFSTKNPAYPI   360
                                 *                  :. . :**.: ..:.* *: :          *   **

sp|Q36736.1|KM11_LEIDO        K-----------------------------------------------------------   92
1PFO:A|PDBID|CHAIN|SEQUENCE   SYTSVFLKDNSVAAVHNKTDYIETTSTEYSKGKINLDHSGAYVAQFEVAWDEVSYDKEGN   420
                              .

sp|Q36736.1|KM11_LEIDO        ------------------------------------------------------------   92
1PFO:A|PDBID|CHAIN|SEQUENCE   EVLTHKTWDGNYQDKTAHYSTVIPLEANARNIRIKARECTGLAWEWWRDVISEYDVPLTN   480

sp|Q36736.1|KM11_LEIDO        --------------------   92
1PFO:A|PDBID|CHAIN|SEQUENCE   NINVSIWGTTLYPGSSITYN   500
```

**Figure S30: Sequence alignment of KMP-11 and Perfringolysin O (PDB ID:1PFO) showed 11.8 % similarity.**

```
CLUSTAL O(1.2.4) multiple sequence alignment


sp|Q36736.1|KM11_LEIDO           ---------MATTYEEFSAKLDRL---DQEFNRKMQ--------E---------------   25
1GWY:A|PDBID|CHAIN|SEQUENCE      ALAGTIIAGASLTFQVLDKVLEELGKVSRKIAVGIDNESGGTWTALNAYFRSGTTDVILP   60
1GWY:B|PDBID|CHAIN|SEQUENCE      ALAGTIIAGASLTFQVLDKVLEELGKVSRKIAVGIDNESGGTWTALNAYFRSGTTDVILP   60
                                           : *:: :.  *:.*   .:::   ::

sp|Q36736.1|KM11_LEIDO           -----QNAKFFADKPDESTLSPEMREHY--------------------------------   48
1GWY:A|PDBID|CHAIN|SEQUENCE      EFVPNTKALLYSGRKDTGPVATGAVAAFAYYMSSGNTLGVMFSVPFDYNWYSNWWDVKIY   120
1GWY:B|PDBID|CHAIN|SEQUENCE      EFVPNTKALLYSGRKDTGPVATGAVAAFAYYMSSGNTLGVMFSVPFDYNWYSNWWDVKIY   120
                                       :* :::.: * . ::      :

sp|Q36736.1|KM11_LEIDO           --------EKFERMIKEHTEKFNKKMHEHSEHFKQKFAELLEQQKAAQYPSK---        92
1GWY:A|PDBID|CHAIN|SEQUENCE      SGKRRADQGMYEDLYYGNPYRGDNGWHEKNLGYGLRMKGIMTSAGEAKMQIKISR        175
1GWY:B|PDBID|CHAIN|SEQUENCE      SGKRRADQGMYEDLYYGNPYRGDNGWHEKNLGYGLRMKGIMTSAGEAKMQIKISR        175
                                           :* :    :  : ::  **:.  :  ::  :: .   *:   *
```

**Figure S31: Sequence alignment of KMP-11 and Sticholysin II (PDB ID:1GWY) showed 26.8% similarity.**

```
sp|Q36736.1|KM11_LEIDO          --MATTYEEFSAK-----------------------------------------------   11
4HSC:X|PDBID|CHAIN|SEQUENCE     MSNKKTFKKYSRVAGLLTAALIIGNLVTANAESNKQNTASTETTTTNEQPKPESSELTTE   60
                                    .*::::*

sp|Q36736.1|KM11_LEIDO          ------------------------------------------------------------   11
4HSC:X|PDBID|CHAIN|SEQUENCE     KAGQKTDDMLNSNDMIKLAPKEMPLESAEKEEKKSEDKKKSEEDHTEEINDKIYSLNYNE   120

sp|Q36736.1|KM11_LEIDO          ----------------------------------------------LDRLDQEFNRKMQE   25
4HSC:X|PDBID|CHAIN|SEQUENCE     LEVLAKNGETIENFVPKEGVKKADKFIVIERKKKNINTTPVDISIIDSVTDRTYPAALQL   180
                                                                               .  *: :   :*

sp|Q36736.1|KM11_LEIDO          QNAKFFADKPDESTLSPE----------MR---------EHYEKFERMIKEHTE------   60
4HSC:X|PDBID|CHAIN|SEQUENCE     ANKGFTENKPDAVVTKRNPQKIHIDLPGMGDKATVEVNDPTYANVSTAIDNLVNQWHDNY   240
                                 *  *  :***  . . :          *            * :..  *.: .:

sp|Q36736.1|KM11_LEIDO          ------------------------------------------------------------   60
4HSC:X|PDBID|CHAIN|SEQUENCE     SGGNTLPARTQYTESMVYSKSQIEAALNVNSKILDGTLGIDFKSISKGEKKVMIAAYKQI   300

sp|Q36736.1|KM11_LEIDO          --------------KF--------------------------------------------   62
4HSC:X|PDBID|CHAIN|SEQUENCE     FYTVSANLPNNPADVFDKSVTFKELQRKGVSNEAPPLFVSNVAYGRTVFVKLETSSKSND   360
                                               *

sp|Q36736.1|KM11_LEIDO          -----NKKMHEHSEHFKQKFAELLEQ----------------------------------   83
4HSC:X|PDBID|CHAIN|SEQUENCE     VEAAFSAALKGTDVKTNGKYSDILENSSFTAVVLGGDAAEHNKVVTKDFDVIRNVIKDNA   420
                                     .  ::  . : : *::::**:

sp|Q36736.1|KM11_LEIDO          ---QKAAQYPSK------------------------------------------------   92
4HSC:X|PDBID|CHAIN|SEQUENCE     TFSRKNPAYPISYTSVFLKNNKIAGVNNRTEYVETTSTEYTSGKINLSHQGAYVAQYEIL   480
                                   :*   ** .

sp|Q36736.1|KM11_LEIDO          ------------------------------------------------------------   92
4HSC:X|PDBID|CHAIN|SEQUENCE     WDEINYDDKGKEVITKRRWDNNWYSKTSPFSTVIPLGANSRNIRIMARECTGLAWEWWRK   540

sp|Q36736.1|KM11_LEIDO          -------------------------------        92
4HSC:X|PDBID|CHAIN|SEQUENCE     VIDERDVKLSKEINVNISGSTLSPYGSITYK        571
```

**Figure S32: Sequence alignment of KMP-11 and Streptolysin O (PDB ID:4HSC) showed 9 % similarity.**

```
sp|Q36736.1|KM11_LEIDO           ------------------------------------------------------------   0
3HVN:A|PDBID|CHAIN|SEQUENCE      GPLGSRKSSHLILSSIVSLALVGVTPLSVLADSKQDINQYFQSLTYEPQEILTNEGEYID   60

sp|Q36736.1|KM11_LEIDO           ------------------------------------------------------------   0
3HVN:A|PDBID|CHAIN|SEQUENCE      NPPATTGMLENGRFVVLRREKKNITNNSADIAVIDAKAANIYPGALLRADQNLLDNNPTL   120

sp|Q36736.1|KM11_LEIDO           ------------------------------------------------------------   0
3HVN:A|PDBID|CHAIN|SEQUENCE      ISIARGDLTLSLNLPGLANGDSHTVVNSPTRSTVRTGVNNLLSKWNNTYAGEYGNTQAEL   180

sp|Q36736.1|KM11_LEIDO           ----------------MATTYEEFSAKLDRLDQEFNR-------------------KMQE   25
3HVN:A|PDBID|CHAIN|SEQUENCE      QYDETMAYSMSQLKTKFGTSFEKIAVPLDINFDAVNSGEKQVQIVNFKQIYYTVSVDEPE   240
                                                 :.*::*:::. **   : .*                    .  *

sp|Q36736.1|KM11_LEIDO           QNAKFFADKPDESTL-----SPE-------------------------------------   43
3HVN:A|PDBID|CHAIN|SEQUENCE      SPSKLFAEGTTVEDLKRNGITDEVPPVYVSSVSYGRSMFIKLETSSRSTQVQAAFKAAIK   300
                                 . :*:**:    . *     : *

sp|Q36736.1|KM11_LEIDO           ----------------------------------MREHYEKFERMIKEHTEKFNKK----   65
3HVN:A|PDBID|CHAIN|SEQUENCE      GVDISGNAEYQDILKNTSFSAYIFGGDAGSAATVVSGNIETLKKIIEEG-ARYGKLNLGV   359
                                                                   :  : *.::::*:*   ::.*

sp|Q36736.1|KM11_LEIDO           ------------------------------------MHEHSEHFKQKFAEL---------   80
3HVN:A|PDBID|CHAIN|SEQUENCE      PISYSTNFVKDNRPAQILSNSEYIETTSTVHNSSALTLDHSGAYVAKYNITWEEVSYNEA   419
                                                                       :**  :  *:

sp|Q36736.1|KM11_LEIDO           -----------------------------------------------------------L   81
3HVN:A|PDBID|CHAIN|SEQUENCE      GEEVWEPKAWDKNGVNLTSHWSETIQIPGNARNLHVNIQECTGLAWEWWRTVYDKDLPLV   479
                                                                                            :

sp|Q36736.1|KM11_LEIDO           EQQKAA-----QYPSK-------------   92
3HVN:A|PDBID|CHAIN|SEQUENCE      GQRKITIWGTTLYPQYADEVIELERPHRD   508
                                  *:* :      **.
```

**Figure S33: Sequence alignment of KMP-11 and Suilysin (PDB ID:3HVN) showed 10.6 % similarity.**

```
CLUSTAL O(1.2.4) multiple sequence alignment


sp|Q36736.1|KM11_LEIDO          ------------------------------------------------------------    0
1XEZ:A|PDBID|CHAIN|SEQUENCE     GAMGSNINEPSGEAADIISQVADSHAIKYYNAADWQAEDNALPSLAELRDLVINQQKRVL    60


sp|Q36736.1|KM11_LEIDO          ------------------------------------------------------------    0
1XEZ:A|PDBID|CHAIN|SEQUENCE     VDFSQISDAEGQAEMQAQFRKAYGVGFANQFIVITEHKGELLFTPFDRTEEIDPALLEAP    120


sp|Q36736.1|KM11_LEIDO          ------------------------------------------------------------    0
1XEZ:A|PDBID|CHAIN|SEQUENCE     RTAALLGASGFASPAPANSETNTLPHVAFYISVNRAISDEECTFNNSWLWKNEKGSRPFC    180


sp|Q36736.1|KM11_LEIDO          ------------------------------------------------MATTYEEFSAKL    12
1XEZ:A|PDBID|CHAIN|SEQUENCE     KDANISLIYRVNLERSLQYGIVGSATPDAKIVRISLDDDSTGAGIHLNDQLGYRQFGASY    240
                                                                                    *.:*.*.

sp|Q36736.1|KM11_LEIDO          DRLDQEFNR---KMQEQNAKFFADKPDE-STLSP--EMREHYEKFERM--------IK--    56
1XEZ:A|PDBID|CHAIN|SEQUENCE     TTLDAYFREWSTDAIAQDYRFVFNASNNKAQILKTFPVDNINEKFERKEVSGFELGVTGG    300
                                  **  *..   .   *: :*. :  :: : :     : :  *****         :.

sp|Q36736.1|KM11_LEIDO          -------------------------EHTEKFNKKM-----HEHSEHF-KQKF---AELLE    82
1XEZ:A|PDBID|CHAIN|SEQUENCE     VEVSGDGPKAKLEARASYTQSRWLTYNTQDYRIERNAKNAQAVSFTWNRQQYATAESLLN    360
                                                          :*:.:. :      :  *  : :*::    .**:

sp|Q36736.1|KM11_LEIDO          QQK----AAQYPSK----------------------------------------------    92
1XEZ:A|PDBID|CHAIN|SEQUENCE     RSTDALWVNTYPVDVNRISPLSYASFVPKMDVIYKASATETGSTDFIIDSSVNIRPIYNG    420
                                :..    .  ** .

sp|Q36736.1|KM11_LEIDO          ------------------------------------------------------------    92
1XEZ:A|PDBID|CHAIN|SEQUENCE     AYKHYYVVGAHQSYHGFEDTPRRRITKSASFTVDWDHPVFTGGRPVNLQLASFNNRCIQV    480


sp|Q36736.1|KM11_LEIDO          ------------------------------------------------------------    92
1XEZ:A|PDBID|CHAIN|SEQUENCE     DAQGRLTANMCDSQQSAQSFIYDQLGRYVSASNTKLCLDGAALDALQPCNQNLTQRWEWR    540


sp|Q36736.1|KM11_LEIDO          ------------------------------------------------------------    92
1XEZ:A|PDBID|CHAIN|SEQUENCE     KGTDELTNVYSGESLGHDKQTGELGLYASSNDAVSLRTITAYTDVFNAQESSPILGYTQG    600


sp|Q36736.1|KM11_LEIDO          ------------------------------------------------------------    92
1XEZ:A|PDBID|CHAIN|SEQUENCE     KMNQQRVGQDNRLYVRAGAAIDALGSASDLLVGGNGGSLSSVDLSGVKSITATSGDFQYG    660


sp|Q36736.1|KM11_LEIDO          ------------------------------------------------------------    92
1XEZ:A|PDBID|CHAIN|SEQUENCE     GQQLVALTFTYQDGRQQTVGSKAYVTNAHEDRFDLPDAAKITQLKIWADDWLVKGVQFDL    720


sp|Q36736.1|KM11_LEIDO          -      92
1XEZ:A|PDBID|CHAIN|SEQUENCE     N      721
```

**Figure S34: Sequence alignment of KMP-11 and Vibrio cholerae cytolysin (PDB ID:1XEZ) showed 7.76% similarity.**

```
sp|Q36736.1|KM11_LEIDO          ------------------------------------------------------------    0
4OWJ:A|PDBID|CHAIN|SEQUENCE     GSAMAHVTLQSLSNNDLCLDVYGENGDKTVAGGSVNGWSCHGSWNQVWGLDKEERYRSRV    60
4OWJ:B|PDBID|CHAIN|SEQUENCE     GSAMAHVTLQSLSNNDLCLDVYGENGDKTVAGGSVNGWSCHGSWNQVWGLDKEERYRSRV    60
4OWJ:C|PDBID|CHAIN|SEQUENCE     GSAMAHVTLQSLSNNDLCLDVYGENGDKTVAGGSVNGWSCHGSWNQVWGLDKEERYRSRV    60
4OWJ:D|PDBID|CHAIN|SEQUENCE     GSAMAHVTLQSLSNNDLCLDVYGENGDKTVAGGSVNGWSCHGSWNQVWGLDKEERYRSRV    60
4OWJ:E|PDBID|CHAIN|SEQUENCE     GSAMAHVTLQSLSNNDLCLDVYGENGDKTVAGGSVNGWSCHGSWNQVWGLDKEERYRSRV    60
4OWJ:F|PDBID|CHAIN|SEQUENCE     GSAMAHVTLQSLSNNDLCLDVYGENGDKTVAGGSVNGWSCHGSWNQVWGLDKEERYRSRV    60
4OWJ:G|PDBID|CHAIN|SEQUENCE     GSAMAHVTLQSLSNNDLCLDVYGENGDKTVAGGSVNGWSCHGSWNQVWGLDKEERYRSRV    60

sp|Q36736.1|KM11_LEIDO          -----------MATTYEEFSAKLDR-----LDQEFNRKMQEQNAKFFADKPD--ESTLSP    42
4OWJ:A|PDBID|CHAIN|SEQUENCE     ASDRCLTVNADKTLTVEQCGANLAQKWYWEGDKLISRYVDGNNTRYLLNIVGGRNVQVTP    120
4OWJ:B|PDBID|CHAIN|SEQUENCE     ASDRCLTVNADKTLTVEQCGANLAQKWYWEGDKLISRYVDGNNTRYLLNIVGGRNVQVTP    120
4OWJ:C|PDBID|CHAIN|SEQUENCE     ASDRCLTVNADKTLTVEQCGANLAQKWYWEGDKLISRYVDGNNTRYLLNIVGGRNVQVTP    120
4OWJ:D|PDBID|CHAIN|SEQUENCE     ASDRCLTVNADKTLTVEQCGANLAQKWYWEGDKLISRYVDGNNTRYLLNIVGGRNVQVTP    120
4OWJ:E|PDBID|CHAIN|SEQUENCE     ASDRCLTVNADKTLTVEQCGANLAQKWYWEGDKLISRYVDGNNTRYLLNIVGGRNVQVTP    120
4OWJ:F|PDBID|CHAIN|SEQUENCE     ASDRCLTVNADKTLTVEQCGANLAQKWYWEGDKLISRYVDGNNTRYLLNIVGGRNVQVTP    120
4OWJ:G|PDBID|CHAIN|SEQUENCE     ASDRCLTVNADKTLTVEQCGANLAQKWYWEGDKLISRYVDGNNTRYLLNIVGGRNVQVTP    120
                                            : * *: .*:* :      *: :.* :: :*:::: :  .  :  ::*

sp|Q36736.1|KM11_LEIDO          EMREHYEKFERMIKEHTEKFNKKMHEHSEHFKQKFAELLEQQKAAQYPSK    92
4OWJ:A|PDBID|CHAIN|SEQUENCE     ENEANQARWK--------------------------PTLQQVKL------    138
4OWJ:B|PDBID|CHAIN|SEQUENCE     ENEANQARWK--------------------------PTLQQVKL------    138
4OWJ:C|PDBID|CHAIN|SEQUENCE     ENEANQARWK--------------------------PTLQQVKL------    138
4OWJ:D|PDBID|CHAIN|SEQUENCE     ENEANQARWK--------------------------PTLQQVKL------    138
4OWJ:E|PDBID|CHAIN|SEQUENCE     ENEANQARWK--------------------------PTLQQVKL------    138
4OWJ:F|PDBID|CHAIN|SEQUENCE     ENEANQARWK--------------------------PTLQQVKL------    138
4OWJ:G|PDBID|CHAIN|SEQUENCE     ENEANQARWK--------------------------PTLQQVKL------    138
                                * . :  :::                            *:* *
```

**Figure S35: Sequence alignment of KMP-11 and Vibrio vulnificus hemolysin (PDB ID:4OWJ) showed 25% similarity.**

#### References

1. Silverman, B. D., Hydrophobic moments of protein structures: Spatially profiling the distribution. *Proceedings of the National Academy of Sciences* **2001,** *98* (9), 4996-5001.
2. Israelachvili, J.; Pashley, R., The hydrophobic interaction is long range, decaying exponentially with distance. *Nature* **1982,** *300* (5890), 341.
3. Sajadi, F.; Rowley, C. N., Simulations of lipid bilayers using the CHARMM36 force field with the TIP3P-FB and TIP4P-FB water models. *PeerJ* **2018,** *6*, e5472.
